# Rpadrino: an R package to access and use PADRINO, an open access database of Integral Projection Models

**DOI:** 10.1101/2022.03.02.482673

**Authors:** Sam C. Levin, Sanne Evers, Tomos Potter, Mayra Peña Guerrero, Dylan Z. Childs, Aldo Compagnoni, Tiffany M. Knight, Roberto Salguero-Gómez

## Abstract

1. Discrete time structured population projection models are an important tool for studying population dynamics. Within this field, Integral Projection Models (IPMs) have become a popular method for studying populations structured by continuously distributed traits *(e.g.* height, weight). Databases of discrete time, discrete state structured population models, for example DATLife (life tables) and COMPADRE & COMADRE (matrix population models), have made quantitative syntheses straightforward to implement. These efforts allow researchers to address questions in both basic and applied ecology and evolutionary biology. There are now over 300 peer-reviewed publications containing IPMs. We describe a novel framework to quickly reconstruct these models for subsequent analyses using *Rpadrino R* package, which serves as an interface to PADRINO, a new database of structured population models.
2. We introduce an R package, *Rpadrino,* which enables users to download, subset, reconstruct, and extend published IPMs. *Rpadrino* makes use of recently created software, *ipmr,* to provide an engine to reconstruct a wide array of IPMs from their symbolic representations and conduct subsequent analyses. *Rpadrino* and *ipmr* are extensively documented to help users learn their usage.
3. *Rpadrino* currently enables users to reconstruct 280 IPMs from 40 publications that describe the demography of 14 animal and 26 plant species. All of these IPMs are tested to ensure they reproduce published estimates. *Rpadrino* provides an interface to augment PADRINO with external data and modify parameter values, creating a platform to extend models beyond their original purpose while retaining full reproducibility.
4. PADRINO and *Rpadrino* provide a toolbox for asking new questions and conducting syntheses with peer-reviewed published IPMs. *Rpadrino* provides a user-friendly interface so researchers do not need to worry about the database structure or syntax, and can focus on their research questions and analyses. Additionally, *Rpadrino* is thoroughly documented, and provides numerous examples of how to perform analyses which are not included in the package’s functionality.

## Introduction

Demography provides an excellent approach to examine the ecology (Crone et al. 2011), evolutionary biology (Metcalf & Pavard 2007), and conservation biology of any species (Doak & Morris 2001). Environmental conditions and biotic interactions influence vital rates *(e.g.* survival, development, reproduction) across the entire life cycle, which then govern its short-term and long-term performance (Caswell 2001). A variety of methods exist for summarizing vital rates into demographic models; discrete-time, structured population models are among the most popular (Crone et al. 2011, Caswell 2001). Indeed, there is a rich history of using such structured population models across a variety of subdisciplines in ecology (*e.g.* Leslie 1945, Caswell 2001, Easterling et al. 2000, Adler et al. 2010, Ellner et al. 2016).

In ecology, matrix projection models (MPMs) are the most widely used structured population model. MPMs divide the population into discrete classes corresponding to some trait value (*e.g.* developmental state, age, or size), and then model the population using vital rates computed for each class. Researchers have also recognized that, for some species, vital rates are best predicted as a function of one or more continuous traits (*e.g.* size, height, mass), rather than as a function of discrete classes (Easterling et al. 2000). Integral projection models (IPMs), which are continuously structured population models, have become an increasingly important tool for ecologists interested in addressing broad biological questions through a demographic lens (Gonzalez et al. 2021). IPMs combine vital rate functions of continuous traits into projection kernels, which describe how the abundance and distribution of trait values in a population change in discrete time (Easterling et al. 2000). IPMs have been used to investigate a variety of topics, such as invasive species spread (*e.g.* Jongejans et al. 2011, Erickson et al. 2017), evolutionary stable strategies (*e.g.* Childs et al. 2004), the effect of climate drivers on population persistence (Salguero-Gómez et al. 2012, Compagnoni et al. 2021a), and linking evolutionary feedbacks to population dynamics (Coulson et al. 2011).

The popularity of MPMs and the desire to conduct quantitative syntheses led to the creation of databases that enable researchers to reproduce and analyze published MPMs en masse (Salguero-Gómez et al. 2014, Salguero-Gómez et al. 2015). There has not been any similar development for IPMs, despite there now being at least 300 publications covering over 380 species (Figure S1, Supplementary Data). Currently, the only way to reproduce these IPMs is to consult the original publication and hope that there is sufficient information in them to fully reproduce the model. This both time consuming and often not possible, as critical details are usually missing (*e.g.* numerical integration rules, range of trait values, SC Levin, personal observation). Given the existing volume of data and its rate of increase, the field of ecology would benefit from a centralized repository that enables researchers to reconstruct and analyze these IPMs. However, storing and reconstructing IPMs present unique challenges that require special attention to enable full use of them.

In order to make full use of the IPM, researchers need, at a minimum, the symbolic representation of the model and the associated parameter values. Existing demographic databases enter the transition values directly, rather than a symbolic matrix and the values associated with the symbols separately. For example, COMPADRE and COMADRE store transition matrices as numeric matrices (sub-matrices corresponding to survival and development (*U*), sexual reproduction (*F*), and asexual reproduction (*C*), and their sum *A*), rather than symbolic matrices with parameter values separately. In general, this limits the variety of subsequent analyses which are possible, because individual matrix elements may be composed of multiple vital rates and this information is lost by storing only the resulting values *(i.e.* the elements of *F* may be comprised of both probability of reproducing and the per-capita number of propagules produced). To avoid this issue for IPMs, we need to store both the functional form of the IPM kernels and vital rate functions, and the associated parameter estimates. We can then use tools that associate the symbols with their values to reproduce the IPMs (*e.g.* metaprogramming and *rlang,* Henry & Wickham 2021). *ipmr* is an *R* package for users to interactively develop their own IPMs from symbolic model representations and parameter estimates, and perform downstream analyses (Levin et al. 2021). *Rpadrino* extends this framework to include *reconstructing* previously published IPMs that are stored in the PADRINO database.

Here, we introduce *Rpadrino. Rpadrino* provides access to PADRINO, which is an open access database of IPMs. Specifically, PADRINO houses symbolic representations of IPMs, their parameter values, and associated metadata to aid users in selecting appropriate models. *Rpadrino* is an R package that enable users to download PADRINO, manage the dataset locally, (optionally) modify, reconstruct, and analyze IPMs from PADRINO. In the following, we describe how to interact with PADRINO using *Rpadrino* and discuss future directions for *Rpadrino* and PADRINO. Importantly, our Supplementary Materials contain two case studies. The first case study demonstrates how to use PADRINO and *Rpadrino* to reconstruct published IPMs, conduct perturbation analyses, compute some life cycle events, and troubleshoot problems. The second case study shows how to use *Rpadrino* and *ipmr* to combine PADRINO IPMs with user-specified IPMs, and then how to use PADRINO data with other databases with BIEN (Maitner et al. 2017) and COMPADRE as examples. The latter is intended to demonstrate the potential for *Rpadrino* in broad, macro-ecological applications. Finally, our supplementary materials also contain a detailed overview of how the PADRINO database, along with the associated assumptions and challenges.

## An introduction to IPMs and PADRINO

First, we provide a brief review of how IPMs are structured. The simplest form of the IPM can be expressed as

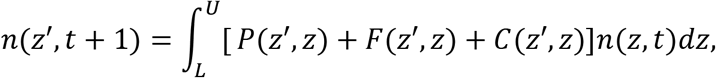

where *n*(*z*’, *t* + 1) and *n*(*z, t*) are the distributions of trait *z* at time *t* + 1 and *t*, *P*(*z’,z*) is a kernel describing survival and development of existing individuals, *F*(*z’, z*) is a kernel describing per-capita sexual reproduction, and *C*(*z’,z*) is a kernel describing per-capita asexual reproduction *(i.e.* clonal reproduction). Each kernel may be comprised of any number of vital rate functions (Ellner, Childs, & Rees 2016). Analytical solutions to the integrals in Eq 1 are not available (Ellner & Rees 2006). Therefore, the integrals are numerically approximated, resulting in a large iteration matrix (typically ranging from 45 × 45 to 5000 × 5000 in dimension, SC Levin unpublished data), and then some quantities of interest are computed (Ellner, Childs & Rees 2016).

Before introducing *Rpadrino,* we also give a very brief overview of PADRINO. PADRINO is an open-access database of integral projection models. PADRINO defines a syntax to symbolically represent IPMs as text strings, and stores the values of those symbols in a separate table. The syntax used is very similar to the mathematical notation of IPMs, and is largely “language-agnostic” *(i.e.* aims to avoid idiosyncrasies of specific programming languages). For example, a survival/growth kernel with the form *P*(*z’,z*) = *s*(*z*) * *G*(*z’,z*) would be `P = s * G` in PADRINO’s syntax. *G*(*z’,z*) = *f_G_*(*z’|μ_g_*(*z*),*σ_G_*) (where *f_G_* denotes a normal probability density function) becomes `G = Norm(mu_g, sd_g)`. This notation should be translatable to many computing languages beyond just *R (e.g.* Python or Julia). Additionally, PADRINO stores extensive metadata to help researchers find IPMs that work for their questions. A more complete description of the database, how IPMs are digitized, and the associated challenges is available in the ESM and the project webpage (Appendix, Tables S1 and S2).

## Rpadrino and ipmr

*Rpadrino* is an *R* package that contains functions for downloading the PADRINO database, data querying and management, modifying existing functional forms and parameter values, and reconstructing models. Model reconstruction is powered by the *ipmr R* package (Levin et al. 2021). While users do not need to know how to use *ipmr* to use *Rpadrino,* the two package are designed to work with and enhance each other. This means that users can combine IPMs reconstructed with *Rpadrino* with IPMs of their own constructed with *ipmr* in a single, coherent analysis (case study 2). Furthermore, users can go from downloading the database to reconstructing IPM objects in as little as 3 function calls. The technical details underlying combining IPM sources are described in *Step 2* below and graphically in Figure 1.

**Figure 1:**
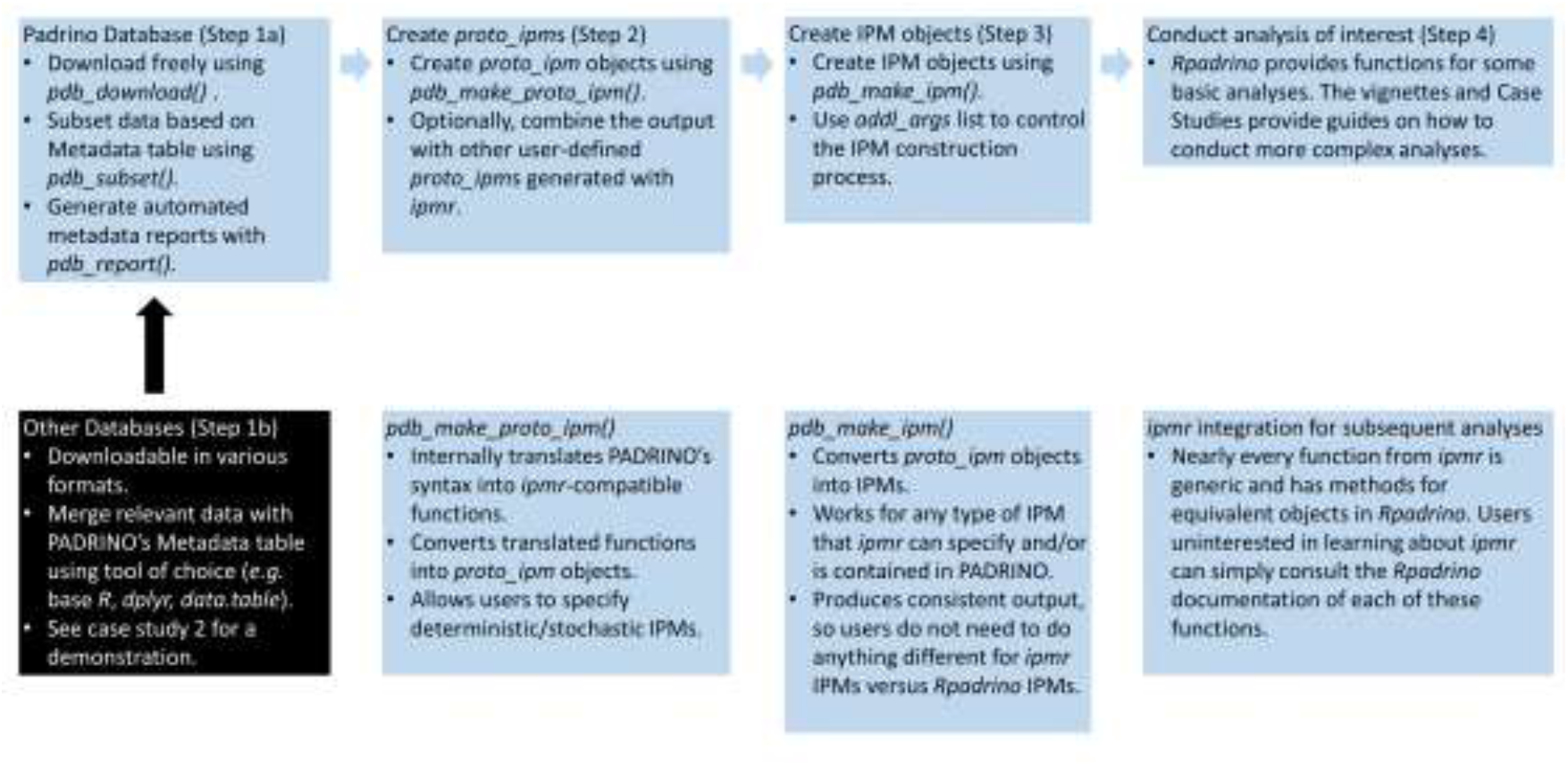
An overview of general workflows with Rpadrino. The first step is to download and subset PADRINO based on the research question at hand (1a). At this point, if the research question calls for it, users may augment the PADRINO object with data from other sources (1b). The extensive metadata table in PADRINO is designed to make this as straightforward as possible. Once the dataset is prepared, users can also generate automated reports on their data subset. After preparing the dataset, users create `proto_ipm` objects using a single function: `pdb_make_proto_ipm()`. This function translates PADRINO’s syntax into ipmr code and then uses ipmr to generate the `proto_ipm` objects. All of these steps take place internally, and so users do not need to understand PADRINO’s syntax or how ipmr works to move forward (2). Once `proto_ipms` are created, users generate IPM objects with a single function, `pdb_make_ipm()`. Examples of how to modify default building settings are in each case study in the supplementary materials and in the function’s documentation (3). Finally, Rpadrino provides some basic analytical machinery, including deterministic and stochastic growth rates and eigenvectors, the ability to make iteration kernels from sub-kernels, and mean kernels (4). Examples of how to conduct more complex analyses are included in the package vignettes and the case studies in the supplementary materials.

The flexibility of IPMs and their broad application across ecology, evolution, and conservation biology mean that there is no fixed set of steps in a workflow using *Rpadrino.* However, there are generally four steps that a researcher must take when using these *Rpadrino.* The first step is to identify studies of interest (Figure 1, Step 1a), and, optionally, augment PADRINO’s metadata with additional information from other sources (*e.g.* environmental data, GBIF, Figure 1, Step 1b). *Rpadrino* represents PADRINO objects as a list of `data. frames` (referred to as tables in subsequent text). *Rpadrino* uses the shared `ipm_id` column across all tables to track information related to each IPM. Therefore, subsetting relies on identifying the correct `ipm_id`’s, and then using those to select the IPMs of interest (Box 1, case study 1 and 2). `data. frames` should be familiar to most *R* users, and the ability to modify them should readily accommodate the range of further analyses that researchers may be interested in. Users may augment any table with additional information corresponding to, for example, spatial or temporal covariates from other open access databases. Furthermore, *Rpadrino* provides numerous access functions for metadata that streamline subsetting (Box 1).

The second step in an *Rpadrino* workflow is to construct a list of `proto_ipm` objects using `pdb_make_proto_ipm()` (Figure 1, Box 1). This function translates PADRINO’s syntax into *ipmr* code, and then builds a `proto_ipm` object for each unique `ipm_id`. For some models, users may choose to create deterministic or stochastic IPMs at this step. *Rpadrino’s* default behavior is to generate deterministic models whenever possible. This behavior encompasses instances where authors generated models with no time or space varying parameters, and where authors included discretely varying environments. The latter can be implemented as deterministic models because all parameter values are known before the IPM is built. IPMs with continuous environmental variation require sampling the environment at each model iteration, usually by sampling from distributions randomly. These are always considered stochastic models. This is also the step where, if needed, users should combine their own `proto_ipm`’s produced by *ipmr* with the `proto_ipm’s` produced by *Rpadrino.*

The third step in an *Rpadrino* workflow is creating IPM objects with `pdb_make_ipm()` (Figure 1, Box 1). `pdb_make_ipm()` uses *ipmrs* `make_ipm()` function to build IPM objects. Users may specify additional options to pass to to `make_ipm()` (*e.g.* normalize the population size to always equal 1, return the vital rate function values as well as the subkernels and population state). The various arguments users can modify are described in the *ipmr* documentation for `make_ipm()`.

The fourth and final step in an *Rpadrino* workflow is to conduct the analyses of interest (Figure 1, Box 1). *Rpadrino* provides functions to extract per-capita growth rates, eigenvectors (Caswell 2001, Ellner, Childs & Rees 2016 Ch. 2, demonstrated in Box 1), assess convergence to asymptotic dynamics (Caswell 2001), compute mean kernels for stochastic IPMs (Ellner, Childs & Rees Ch 7), and modify existing IPMs with new parameter values and functional forms. Additionally, the documentation on the Rpadrino website and the Supplementary Materials for this paper contain details on how to conduct more complicated analyses with IPM objects *(e.g.* perturbation analyses (Ellner, Childs & Rees 2016 Ch 4), size at death calculations (Metcalf et al. 2009)). The package documentation and the recent publication describing *ipmr* also contain code demonstrating analyses on single IPM objects (Levin et al. 2021). These can be extended via the apply family of functions.

**Box 1:**
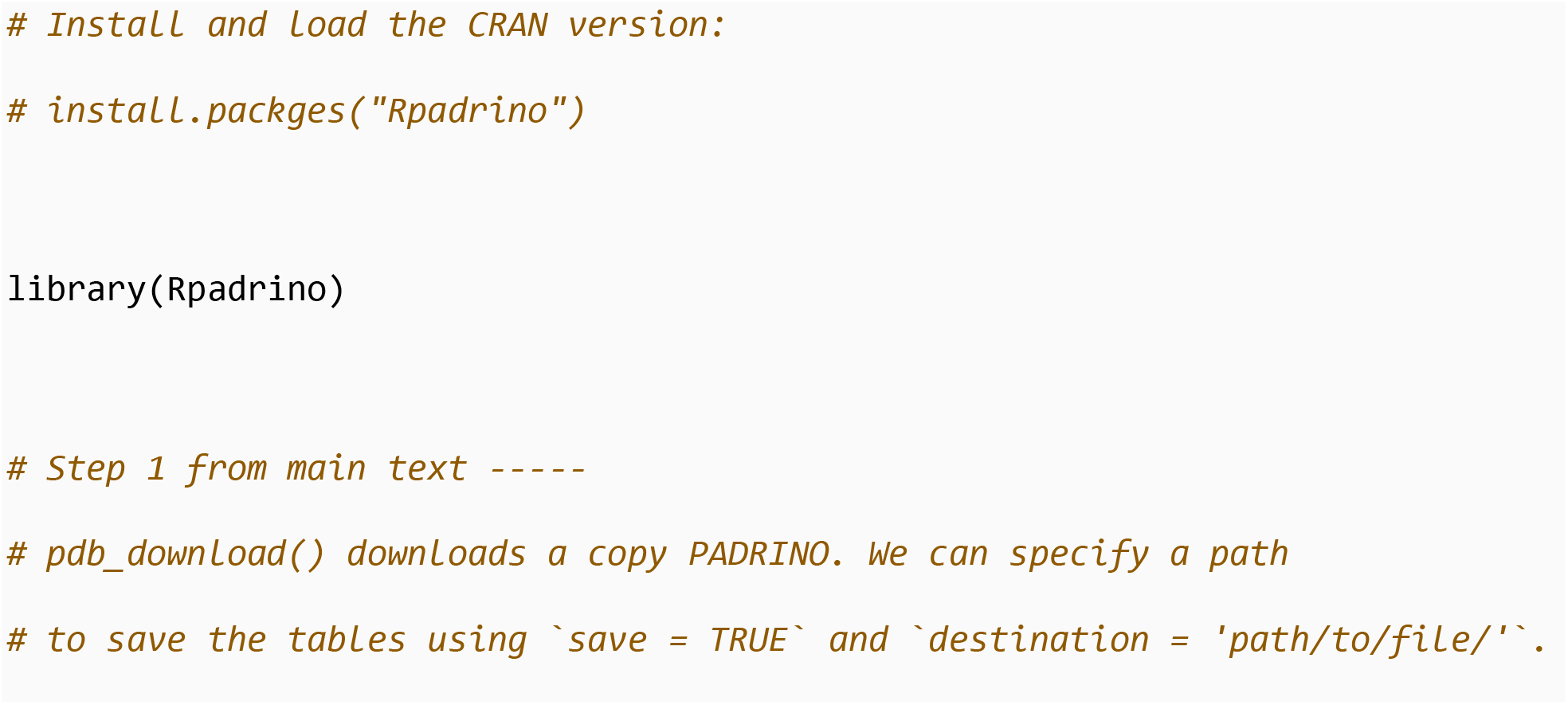
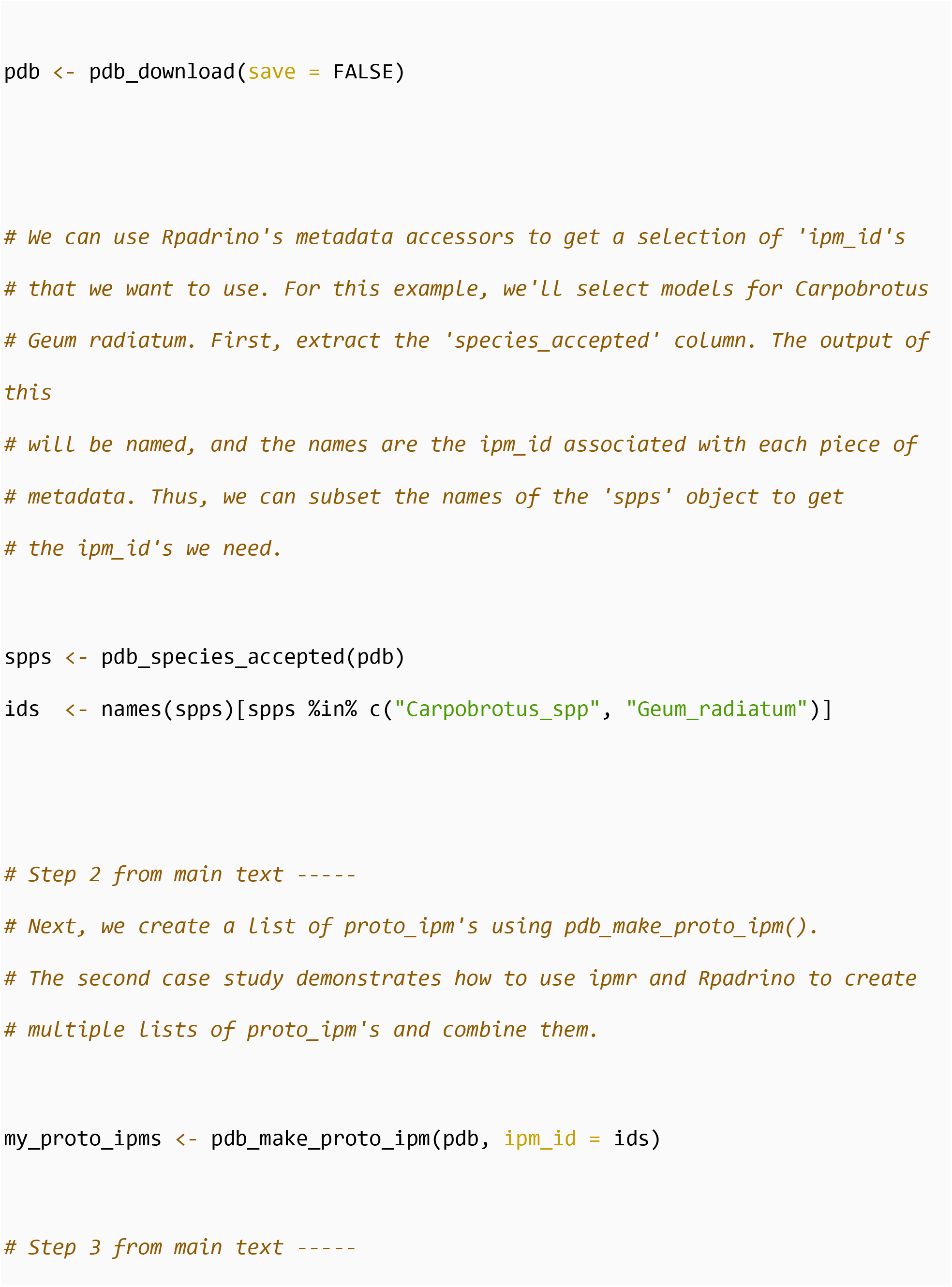
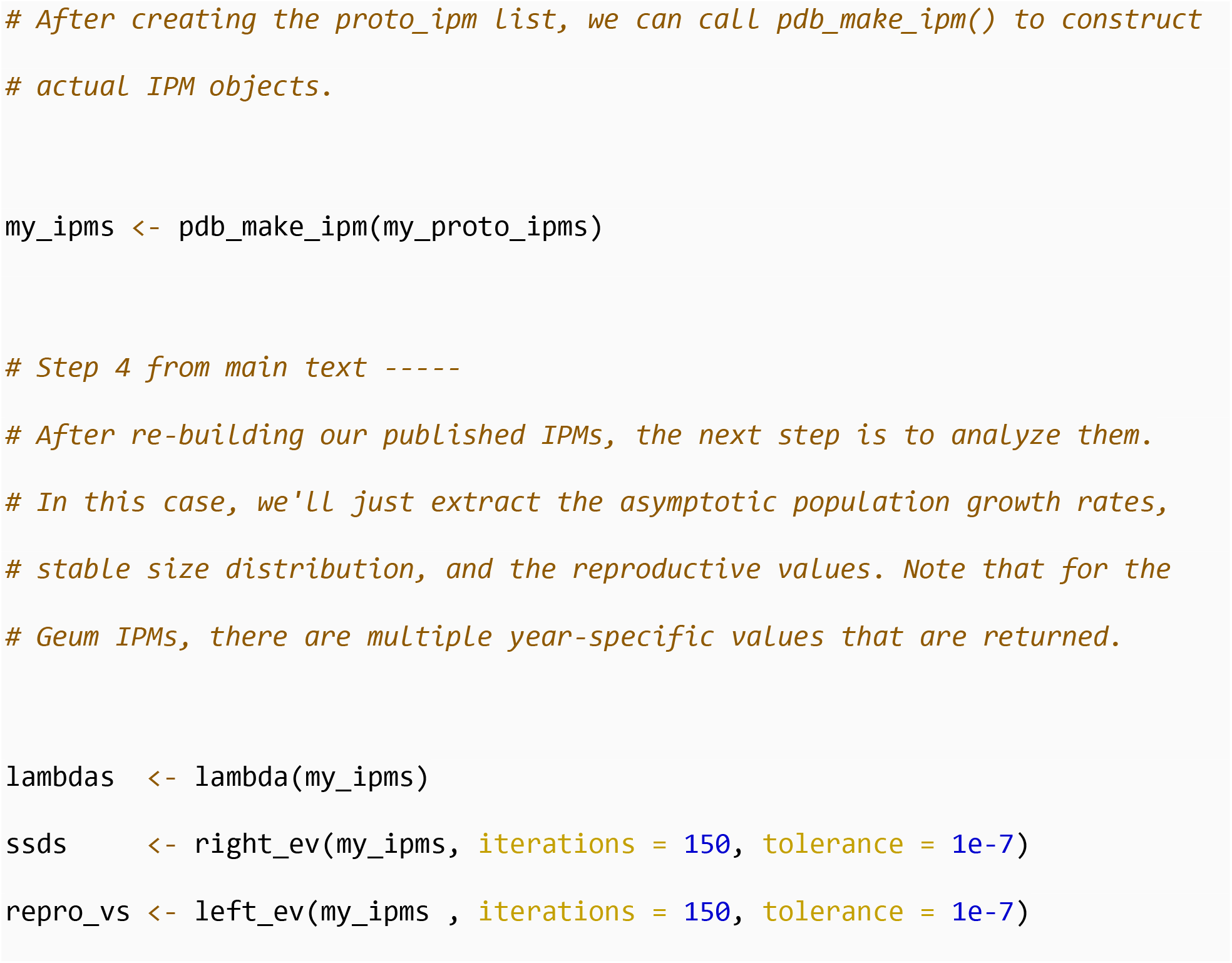
*An example of a simple analysis workflow using* Rpadrino *. The first step in using* Rpadrino *is to install and load the package. After that, we can use* RPadrino *to download* PADRINO *and, optionally, save it locally on our computer. Once the data are downloaded, we can make use of* Rpadrino*’s metadata accessor functions to quickly select models that meet our criteria (step 1). The concept of the* `ipm_id` *is explained in greater detail in the Appendix of this manuscript. The next step is to use these* `ipm_id`*’s to create a list of* `proto_ipm`*’s using* `pdb_make_proto_ipm()` *(step 2). After this step, we can create actual IPM objects using* `pdb_make_ipm()` *(step 3). Once IPM objects are created, the following steps are according to the demands of the research question. In this case, asymptotic population growth rates, stable size distributions, and reproductive values are extracted (step 4). Note that since the* Geum radiatum *model includes a number of year-specific estimates, multiple values are generated for each quantity we want to extract. The concise representation and reconstruction of models such as this is powered by* ipmr*’s parameter set index notation, which is described in greater detail* here. *However, users do not need to be familiar with this notation unless they wish to modify the IPM in question (see case study 1 for an example of modifying* PADRINO *IPMs with* Rpadrino).

## Challenges

There are numerous challenges associated with digitizing and reproducing published IPMs. The digitization side is discussed in Appendix 1, and so is not treated further here.

Important challenges remain in the reconstruction of IPMs. Semi- or non-parametric models may be used to generate IPMs whose functional form is not known *a priori.* We have not yet developed a general syntax for representing these models in PADRINO, though work is ongoing. Additionally, *ipmr* is not yet able to handle two-sex models (*e.g.* Stubberud et al. 2019), time-lagged models (*e.g.* Rose et al. 2005), or periodic models (*e.g.* Letcher et al. 2014). These types of IPMs do not yet represent a substantial portion of the literature. Nonetheless, it is our intention to continue developing functionality to accommodate them in future releases of *Rpadrino, ipmr,* and PADRINO.

## Opportunities and Future Directions

PADRINO is designed to be compatible with other COMPADRE and COMADRE (and others too), and *Rpadrino* is intended to streamline this interoperability. PADRINO’s metadata table closely resembles that of COMPADRE and COMADRE (Salguero-Gómez et al. 2015, Table S2). This means that users who are familiar with these matrix population model databases should find a smooth transition to working with PADRINO. Furthermore, we provide a comprehensive set of metadata that allows users to match information in PADRINO with other databases (*e.g.* gridded climate data, functional trait information, phylogenetic information). These features will enable researchers to carry out more detailed and comprehensive analyses at various spatial, temporal, and phylogenetic scales.

*Rpadrino* presents unique opportunities for synthesis in both theoretical and applied contexts. The expanded range of phylogenetic and geographical coverage can be used in conjunction with other demographic databases (*e.g.* COM(P)ADRE (Salguero-Gómez et al. 2015, Salguero-Gómez et al. 2016), popler (Compagnoni et al. 2019), DatLife (DatLife 2021)) to power larger scale syntheses than were possible before (*e.g.* Compagnoni et al. 2021b). For example, one could use IPMs from PADRINO and matrix population models from COMPADRE and COMADRE to create life tables (Jones et al. 2021), which could then be combined with life tables from DATLife for further analysis (*e.g.* Jones et al. 2014). The intermediate life table conversion steps may not be necessary, as many of the same life history traits and population level parameters may be calculated from all of these models (Caswell 2001, Ellner, Childs & Rees 2016). Furthermore, recent publications combine biotic and abiotic interactions into demographic models providing a robust theoretical toolbox for exploring species responses to environmental drivers such as climate change (*e.g.* Simmonds et al. 2020, Abrego et al. 2021). *Rpadrino* also provides functionality to modify parameter values and functional forms of the IPMs it stores, giving theoreticians a wide array of realistic life histories to experiment with. These examples are far from an exhaustive list, but hopefully demonstrates the potential for this new tool in demography, ecology, and evolutionary biology.

**Table 1:**
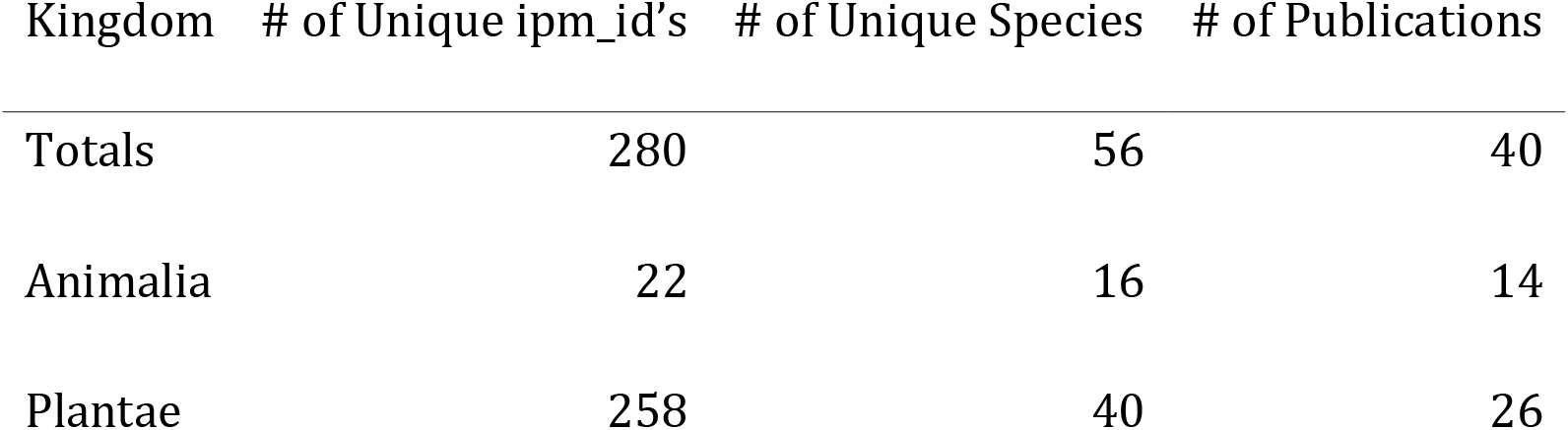
Taxonomic representation of IPMs accesible via Rpadrino. These numbers represent the number of models that are error checked and accurately reproduce the published IPM (see ‘Data Validation’ in the Appendix for more details). Models that are partially entered or still contain errors are not considered here. We are in the process of correcting them and/or retrieving additional information from the authors. See Appendix for details.

| Kingdom | # of Unique ipm_id's | # of Unique Species | # of Publications |
| --- | --- | --- | --- |
| Totals | 280 | 56 | 40 |
| Animalia | 22 | 16 | 14 |
| Plantae | 258 | 40 | 26 |

## Supporting information

Case Study 1

Case Study 2

Electronic Supplementary Info

## Acknowledgments

Funding: R.SG. was supported by a NERC Independent Research Fellowship (NE/M018458/1). SCL, AC, SE, and TMK were funded by the Alexander von Humboldt Foundation in the framework of the Alexander von Humboldt Professorship of TM Knight endowed by the German Federal Ministry of Education and Research.

## Author Contributions

RSG, TMK, DZC, AC, and SCL designed the database. SE, TP, MP-G and SCL collected the data. SCL designed *ipmr, Rpadrino,* and *pdbDigitUtils* with contributions from all authors, and SCL implemented the packages. SCL wrote the first draft of the manuscript and all authors provided comments.

