## Supplementary material for "Rpadrino: an R package to access and use PADRINO, an open access database of Integral Projection Models": Case Study 1

### Stand-alone analyses with PADRINO and Rpadrino

**Mean lifetime recruit production** 16

**Troubleshooting and Modifying IPMs** 21

**Vital rate and parameter level analyses** 23

**Recap** 38

**Citations** 38

### PADRINO and Rpadrino

We have created *Rpadrino* to streamline the process of interacting with PADRINO from *R*. The goal of *Rpadrino* is data management and model construction - not necessarily to do analyses for you. Thus, there is still some minimum amount of programming knowledge required to use it. It will also be helpful to understand how *ipmr*, the engine that powers model reconstruction, creates model objects and the things that it returns. *ipmr* is extensively documented here, and reading at least the introduction to that will certainly help understand the code that follows here. Eventually, we plan to create the *ipmtools* package, which will house functions designed to work with the IPMs stored in PADRINO to conduct more extensive analyses (*e.g.* perturbations, LTREs, life history traits).

### Usage

This case study makes use of *dplyr* to help with data transformation. If you do not already have it, install it (and *Rpadrino*) with:

```
install.packages(c("dplyr", "Rpadrino"))
```

We will show how to compute sensitivity and elasticity for simple models, and then derive some demographic quantities from models housed in PADRINO. The first step is to identify models that have the information we want. Sensitivity and elasticity computations will make use of functions contained in *ipmr*, and we will define a couple of our own to help tie it all together.

After perturbation analyses, we will compute the mean lifetime output of recruits as a function of initial size $z_0$, $\bar{r}(z_0)$. This is defined as $\bar{r}(z_0) = eFN$, where $F$ represents a the fecundity kernel, and $N$ is the fundamental operator. The fundamental operator can be thought of as the expected amount of time spent in any state $z'$ prior to death given an initial state $z_0$ (Caswell 2001), and is computed as $N = (I - P)^{-1}$ (where $P$ is the survival/growth kernel, and $I$ is an identity operator such that $IP = PI = P$).

Finally, we will compute mean size at death ($\bar{\omega}(z_0) = (\mathbf{i} \circ (1 - s))N$), and the size at death kernel ($\Omega(z', z_0) = (1 - s(z'))N(z', z_0)$. Thus, we need models that contain information on survival and growth, and sexual reproduction. To keep things simpler, we will restrict ourselves to simple IPMs.

> NB: The above formulae are from Ellner, Childs, & Rees (2016), Table 3.3. Their derivations are described in detail in Chapter 3 of the book.

#### Subsetting using Metadata and other tables

We can find simple IPMs in PADRINO using a combination of *dplyr* (Wickham et al. 2021) and *Rpadrino* code. *Rpadrino* provides the `pdb_subset()` function. `pdb_subset()` currently only takes `ipm_ids` that we want to keep. The functionality will get expanded, but it's surprisingly complicated to manage that in a user-friendly interface (see the PADRINO explorer app for additional help). This means that we have to work out which `ipm_ids` correspond to the models we want, and then pass those to `pdb_subset()`.

It is a good idea to consult the table guide so that you are familiar with table and variable names, and what information they provide. This case study will introduce some of the variables and tables, but it does not cover them all!

```
library(dplyr)
```

```
## Warning: package 'dplyr' was built under R version 4.1.1
```

```
library(Rpadrino)

pdb <- pdb_download(save = FALSE)

# Simple models only make use of 1 trait/state variable. Therefore, if a model
# has more than 1, it is, by definition, not a simple model. The code below
# calculates the number of traits per "ipm_id", and then filters out those
# that have more than 1 trait. The final piece with the square brackets makes sure
# the final result is a character vector containing only ipm_id's, rather than
# a data.frame

simple_mod_ind <- pdb$StateVariables %>%
  group_by(ipm_id) %>%
  summarise(N = n()) %>%
  filter(N < 2) %>%
  .[ , 1, drop = TRUE]
```

```r
simple_pdb <- pdb_subset(pdb, simple_mod_ind)
```

We have quite a few to choose from! However, a number of these may be stochastic and/or density-dependent models. We will want to get rid of those too, as sensitivity analyses can be trickier and more time consuming for them. This process will use the `Metadata`, `EnvironmentalVariables`, and `ParSetIndices` tables to find those.

```r
# The first piece of stoch_ind examines the EnvironmentalVariables table. This
# contains information on IPMs that include continuous environmental variation.
# Rpadrino treats these as stochastic by default, because  PADRINO almost
# always uses random number generators to sample the distributions of environmental
# values. Therefore, there is not really a way to sample these in a way that makes
# them deterministic while still preserving the published model.

stoch_ind <- unique(simple_pdb$EnvironmentalVariables$ipm_id)

# The second piece of stoch_ind examines the ParSetIndices table. This table
# describes discrete environmental variation. These models do not have to be
# stochastic, but they will make the analysis a bit more complicated, so we are
# going to drop those for now.

stoch_ind <- c(stoch_ind, unique(simple_pdb$ParSetIndices$ipm_id))

# The final piece of stoch_ind checks for density dependence. This information
# is stored in the 'has_dd' column of the Metadata table.

stoch_ind <- c(stoch_ind,
               unique(simple_pdb$Metadata$ipm_id[simple_pdb$Metadata$has_dd]))

det_pdb <- pdb_subset(simple_pdb, setdiff(simple_pdb$Metadata$ipm_id,
                                          stoch_ind))
```

For simple models in PADRINO, the kernels are pretty consistently named with respect to the broader IPM literature: `P` denotes survival and growth, `F` denotes sexual reproduction, and `C` denotes asexual reproduction. The `IpmKernels` table stores the names, functional forms of the kernels, as well as other information needed to implement them. We can use the kernel names column, `kernel_id`, to do a quick sanity check to make sure our subsetting produced only these kernels like so:

```r
unique(det_pdb$IpmKernels$kernel_id)
```

```
## [1] "P" "F"
```

Great, all `P`s and `F`s! Finally, we are going to make sure we only have one IPM per species in our analysis. We can do this using the `Metadata` table. This is certainly not required for any analyses, just to keep things tractable for now.

```r
keep_ind <- det_pdb$Metadata$ipm_id[!duplicated(det_pdb$Metadata$species_accepted)]

my_pdb <- pdb_subset(det_pdb, keep_ind)
```

#### Re-building IPMs

Now that we have our data subsetted, we can start making IPMs. The first step is always to create `proto_ipm` objects. These are an intermediate step between the database and a set of usable kernels. Because *ipmr* also uses these as an intermediate step, we can combine models from PADRINO with ones that we create ourselves. There is an example of this in the second case study.

##### Under the hood

If you're interested in how PADRINO actually stores IPMs, and specifically the expressions that comprise them, keep reading. If not, skip to the next heading.

Lets have a look at how PADRINO stores kernels, vital rates, and parameters. These are in the `IpmKernels`, `VitalRateExpr`, and `ParameterValuess` tables, respectively.

```
head(my_pdb$IpmKernels[ , 1:3])
```

```
##    ipm_id kernel_id                                             formula
## 43 aaaa34         P                                 P = s * g * d_lnsize
## 44 aaaa34         F                      F = r * fn * pE * d  * d_lnsize
## 47 aaaa36         P                                 P = s * g * d_lnsize
## 48 aaaa36         F                      F = r * fn * pE * d  * d_lnsize
## 92 aaa144         P                                   P = s * g * d_size
## 93 aaa144         F F = rep_p * es_p * sdl_s *n_infl * n_fl * n_seed * d_size
```

```
head(my_pdb$VitalRateExpr[ , 1:3])
```

```
##     ipm_id demographic_parameter                                  formula
## 147 aaaa34              Survival s = 1/(1+exp(-(s_b + s_m * lnsize_1)))
## 148 aaaa34                Growth                  g = Norm(g_mean, g_var)
## 149 aaaa34                Growth            g_mean = g_b + g_m * lnsize_1
## 150 aaaa34                Growth   g_var = sqrt(gv_b + gv_m * lnsize_1)
## 151 aaaa34             Fecundity r = 1/(1+exp(-(r_b + r_m * lnsize_1)))
## 152 aaaa34             Fecundity       fn = exp(fn_b + fn_m * lnsize_1)
```

```
head(my_pdb$ParameterValues)
```

```
##     ipm_id demographic_parameter state_variable parameter_name parameter_value
## 619 aaaa34              Survival         lnsize            s_b      -0.5612335
## 620 aaaa34              Survival         lnsize            s_m       0.4628431
## 621 aaaa34                Growth         lnsize            g_b       1.1088198
## 622 aaaa34                Growth         lnsize            g_m       0.5148672
## 623 aaaa34                Growth         lnsize           gv_b       0.9504887
## 624 aaaa34                Growth         lnsize           gv_m       0.0000000
```

We can see that the kernels and vital rate expressions are all defined symbolically, and the parameter values are stored elsewhere. This helps us reuse parameters that appear in multiple expressions without re-typing them, reducing the risk of errors. Additionally, it'll make it easier for us to modify parameter values, vital rate expressions, and kernel formulae if we want to. However, the syntax in the tables is probably not the easiest to work with directly. Therefore, *Rpadrino* provides the `pdb_make_proto_ipm()` function. This takes a `pdb` object and produces a list of `proto_ipm`s. In the chunk after this one, we will see that it translates the syntax in `IpmKernels` and `VitalRateExpr` into usable `R` code. There are additional options that we can pass to this, but we will ignore those for now, and just focus on creating and understanding what the outputs are.

##### Creating the proto_ipm list

The following line generates a set of `proto_ipm`'s for the species in our subsetted database:

```
simple_det_list <- pdb_make_proto_ipm(my_pdb)
```

```
## 'ipm_id' aaa310 has the following notes that require your attention:
## aaa310: 'Geo and time info retrieved from COMPADRE (v.X.X.X.4)'

## 'ipm_id' aaa323 has the following notes that require your attention:
## aaa323: 'Simulated demographic data derived from Nicole J Ecol 2011'

## 'ipm_id' aaa326 has the following notes that require your attention:
## aaa326: 'Demographic data from Metcalf Funct Ecol 2006'

## 'ipm_id' aaa385 has the following notes that require your attention:
## aaa385: 'Same data as AAA385. State variable Height (Cm)'

## 'ipm_id' ddddd3 has the following notes that require your attention:
## ddddd3: 'Frankenstein IPM'

## 'ipm_id' ddddd4 has the following notes that require your attention:
## ddddd4: 'assumes mean surface temp of 10.34 °C, and constant survival probability of
## large pike'

## 'ipm_id' dddd24 has the following notes that require your attention:
## dddd24: 'Assumes no external recruitment'

## 'ipm_id' dddd26 has the following notes that require your attention:
## dddd26: '1 ipm digitized, additional ipms taking into account dispersal still
## possible to digitize'

## 'ipm_id' dddd30 has the following notes that require your attention:
## dddd30: 'Frankenstein IPM'

## 'ipm_id' dddd37 has the following notes that require your attention:
## dddd37: 'MS contains 2 det and 2 stoch IPMs, only 1 det inlcuded here'

## 'ipm_id' dddd39 has the following notes that require your attention:
## dddd39: 'Only deterministic model included here: assumes precipitation = 104mm'

## 'ipm_id' dddd40 has the following notes that require your attention:
## dddd40: 'DEB-IPM - these vital rates assume NO shrinking. 1 model digitized: assumes
## that scaled functional response E_Y = 0.65, with var(E_Y) = 0.1 - see paper for
## details'

## 'ipm_id' dddd41 has the following notes that require your attention:
## dddd41: 'DEB-IPM - these vital rates shrinking IS possible.  1 model digitized:
## assumes that scaled functional response E_Y = 0.65, with var(E_Y) = 0.1 - see paper
## for details'
```

First, we note that the building process threw out a few messages. The first is that the coordinates and duration information come from COMPADRE, not necessarily the original publication. This is not really alarming - COMPADRE is pretty trustworthy. The next few are related to demographic data sources and GPS location. "Frankenstein IPM" refers to a situation where some vital rates are measured directly from demographic data the authors collected, while other vital rates were retrieved from the literature (*i.e.* the object is cobbled together from disparate sources, Shelley 1818). Again, not necessarily alarming, though we'd want to know that if our study question required that all vital rates come from one place (*e.g.* matching environmental conditions to demographic performance). The next few tell us that the publication actually contains more IPMs, but that our PADRINO digitization team hasn't finished entering all of them yet. And finally, there is a note about the assumptions contained by the model.

we will inspect a couple of the objects in this list to get a feel for what a `proto_ipm` contains:

```
simple_det_list
```

```
## This list of 'proto_ipm's contains the following species:
## Poa alsodes
```

```
## Poa sylvestris
## Aeonium haworthii
## Cotyledon orbiculata
## Aconitum noveboracense
## Dracocephalum austriacum
## Cirsium arvense
## Lonicera maackii
## Mimulus cardinalis
## Reynoutria japonica
## Carpobrotus spp
## Crocodylus niloticus
## Esox lucius
## Sisturus catenatus catenatus
## Ovis aries
## Oncorhynchus clarkii
## Tridacna maxima
## Gadus morhua
## Podarcis lilfordi
## Nerodia sipedon
## Ostrea edulis
## Dipsastraea favus
## Platygyra lamellina
## Ficedula hypoleuca
## Testudo graeca
## Manta alfredi
## Rhizoglyphus robini
##
## You can inspect each model by printing it individually.
```

```
simple_det_list$aaaa34
```

```
## A simple, density independent, deterministic proto_ipm with 2 kernels defined:
##  P, F
##
## Kernel formulae:
##
## P: s * g
## F: r * fn * pE * d
##
## Vital rates:
##
## s: 1/(1 + exp(-(s_b + s_m * lnsize_1)))
## g_mean: g_b + g_m * lnsize_1
## g_var: sqrt(gv_b + gv_m * lnsize_1)
## g: stats::dnorm(lnsize_2, g_mean, g_var)
## r: 1/(1 + exp(-(r_b + r_m * lnsize_1)))
## fn: exp(fn_b + fn_m * lnsize_1)
## d: stats::dexp(lnsize_2, 1/d_mean)
##
## Parameter names:
##
##  [1] "s_b"    "s_m"    "g_b"    "g_m"    "gv_b"   "gv_m"   "r_b"    "r_m"
##  [9] "fn_b"   "fn_m"   "t_r"    "d_mean" "pE"
##
## All parameters in vital rate expressions found in 'data_list':  TRUE
```

```
##
## Domains for state variables:
##
## lnsize: lower_bound = 0, upper_bound = 5, n_meshpoints = 500
##
## Population states defined:
##
## n_lnsize: Pre-defined population state.
##
## Internally generated model iteration procedure:
##
## n_lnsize_t_1: right_mult(kernel = P, vectr = n_lnsize_t) + right_mult(kernel = F,
##     vectr = n_lnsize_t)
```

```
simple_det_list$dddd30
```

```
## A simple, density independent, deterministic proto_ipm with 2 kernels defined:
##  P, F
##
## Kernel formulae:
##
## P: s * g
## F: s * r * pg * 0.5 * d
##
## Vital rates:
##
## s: 1/(1 + exp(-(aS + bS * svl_1 + cS * svl_1^2)))
## muG: svl_1 + (Linf - svl_1) * (1 - exp(-k * tg))
## g: stats::dnorm(svl_2, muG, sigmaG)
## r: exp(aR + bR * svl_1)
## pg: ifelse(svl_1 < svlM, 0, 1)
## d: stats::dnorm(svl_2, muD, sigmaD)
##
## Parameter names:
##
##  [1] "aS"     "bS"     "cS"     "Linf"   "k"      "tg"     "sigmaG" "aR"
##  [9] "bR"     "svlM"   "muD"    "sigmaD"
##
## All parameters in vital rate expressions found in 'data_list':  TRUE
##
## Domains for state variables:
##
## svl: lower_bound = 120, upper_bound = 1200, n_meshpoints = 1000
##
## Population states defined:
##
## n_svl: Pre-defined population state.
##
## Internally generated model iteration procedure:
##
## n_svl_t_1: right_mult(kernel = P, vectr = n_svl_t) + right_mult(kernel = F,
##     vectr = n_svl_t)
```

We can see that *Rpadrino* has translated PADRINO's syntax into a set of *R* expressions that correspond to the vital rate functions and sub-kernel functional forms, as well as checked that the model can be implemented with the parameter values that are present in PADRINO. Finally, it has generated the model iteration expression, which shows how the sub-kernels interact with each trait distribution at time $t$ to produce new trait distributions at time $t+1$. We can now build the actual IPM objects. We will also check for convergence to asymptotic dynamics using the `is_conv_to_asymptotic` function.

```
all_ipms   <- pdb_make_ipm(simple_det_list)

check_conv <- is_conv_to_asymptotic(all_ipms)
```

```
## The following IPMs did not converge: aaa310, aaa341, ccccc1, ddddd3, ddddd5,
## dddd10, dddd24, dddd26, dddd30, dddd33, dddd35, dddd36, dddd37, dddd39,
## dddd40, dddd41
```

```
check_conv
```

```
## [1] FALSE
```

We can see that a few of these need more than the default number of iterations to converge to asymptotic dynamics. All $\lambda$ values are computed via iteration, rather than computing eigenvalues. Since we need correct $\lambda$ values to compute elasticity, we will need to re-run those models until they converge (or at least come very close to convergence). `pdb_make_ipm()` contains the `addl_args` argument that tells the function how to deviate from the default behavior of `ipmr::make_ipm()`. It's accepts nested lists with the following format:

```
list(<ipm_id_1> = list(<make_ipm_arg_name_1> = <XXX>,
                       <make_ipm_arg_name_2> = <YYY>),
     <ipm_id_2> = list(<make_ipm_arg_name_1> = <XXX>,
                       <make_ipm_arg_name_5> = <ZZZ>))
```

We replace the values in `<>` with the actual `ipm_id`s, argument names, and values we want them to have. We can do this many models a bit more concisely:

```
# Create an empty list with names that correspond to ipm_id's that we want to add
# additional iterations for.

ind_conv <- c(paste0("aaa", c(310,341)),
              paste0("ccccc", 1),
              paste0("ddddd", c(3,5)),
              paste0("dddd", c(10, 24, 26, 30, 33, 35, 36, 37, 39, 40, 41)))

# Next, we set create an entry in each list with iterations = <some number>
# we will use 250 for this example. We need to set the names of the list to be
# the ipm_id's, so that pdb_make_ipm() knows which models to use the additional
# arguments with.

arg_list <- lapply(ind_conv,
                   function(x, n_iter) list(iterations = n_iter),
                   n_iter = 250) %>%
  setNames(ind_conv)

new_ipms <- pdb_make_ipm(simple_det_list, addl_args = arg_list)

# Check for convergence out to 5 digits. This should be close enough for what
# we want to do.

check_conv <- is_conv_to_asymptotic(new_ipms, tolerance = 1e-5)

# We can also plot the lambda time series using conv_plot methods for pdb_ipms.
```

```r
# The last two models may not have converged
par(mfrow = c(4, 2))

conv_plot(new_ipms)
```

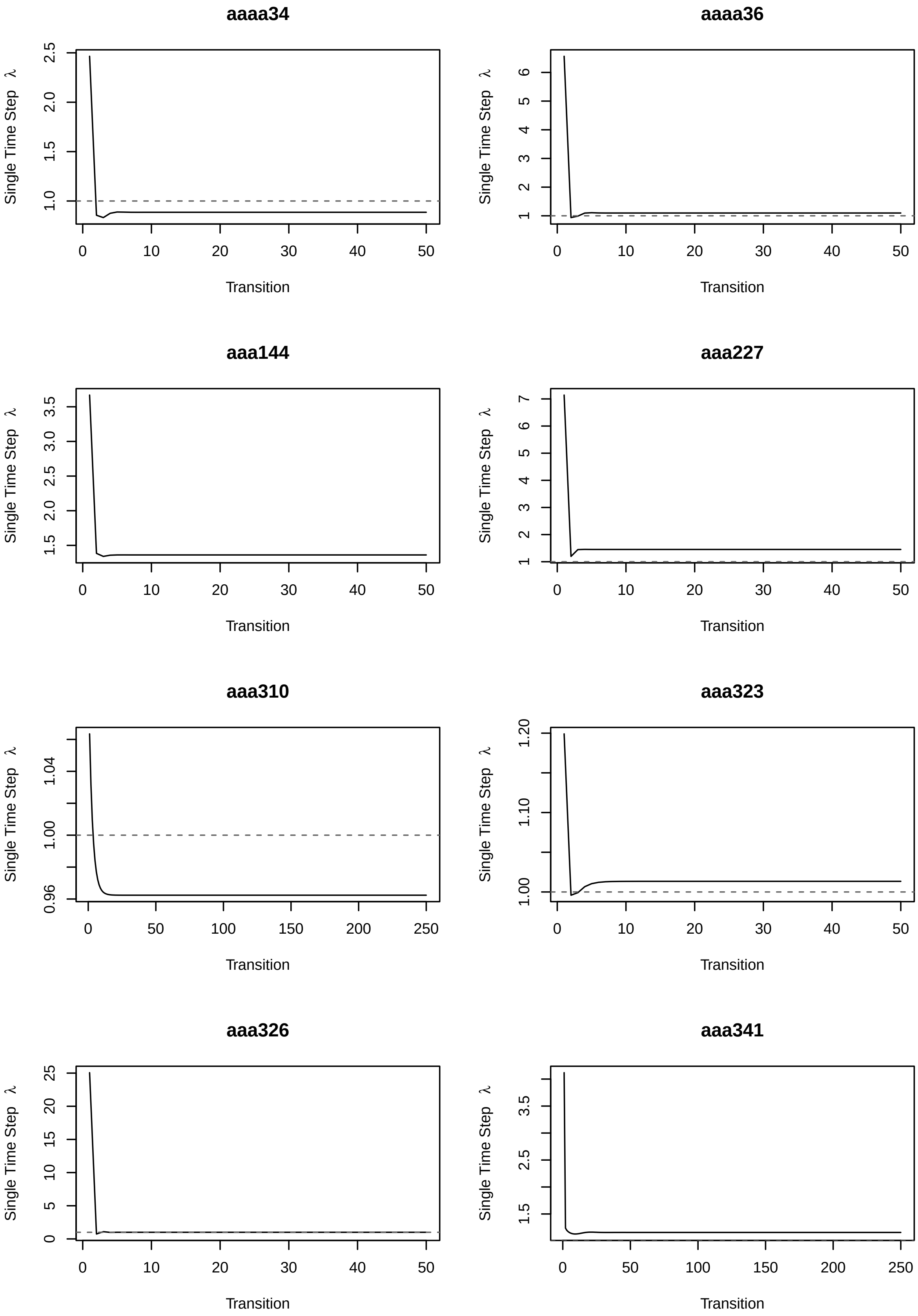
aaaa34
aaaa36
aaa144
aaa227
aaa310
aaa323
aaa326
aaa341
Single Time Step λ
Transition

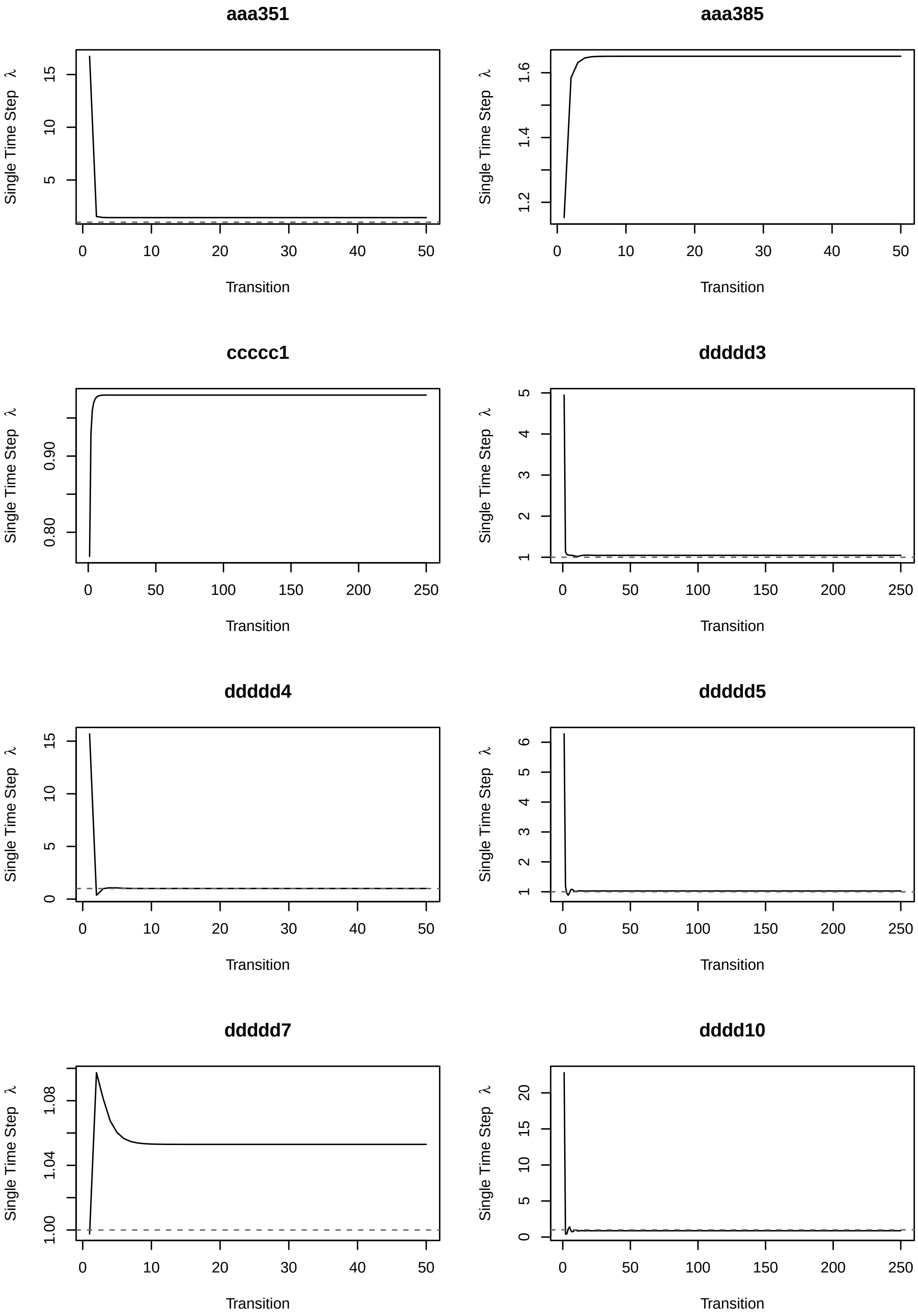
aaa351
aaa385
ccccc1
ddddd3
ddddd4
ddddd5
ddddd7
dddd10
Single Time Step λ
Transition

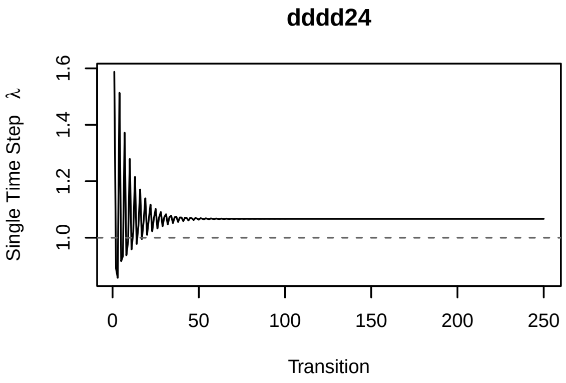
dddd24
Single Time Step λ
1.0 1.2 1.4 1.6
0 50 100 150 200 250
Transition

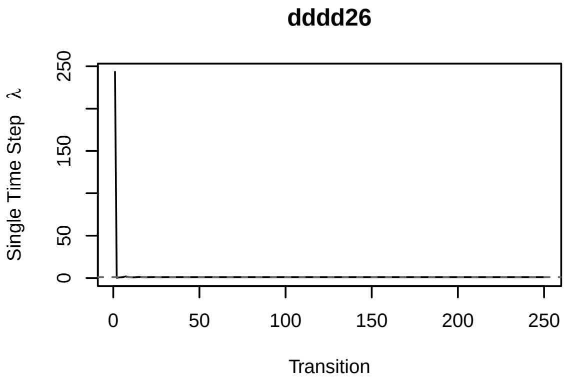
dddd26
Single Time Step λ
0 50 150 250
0 50 100 150 200 250
Transition

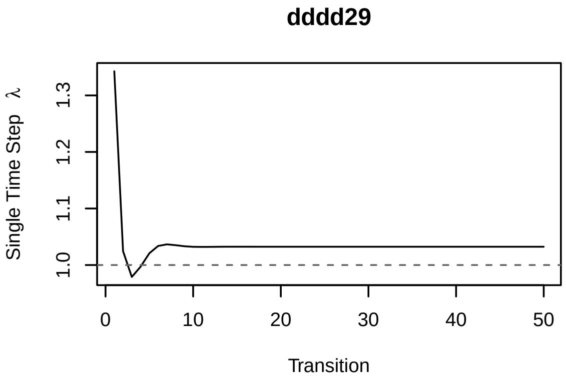
dddd29
Single Time Step λ
1.0 1.1 1.2 1.3
0 10 20 30 40 50
Transition

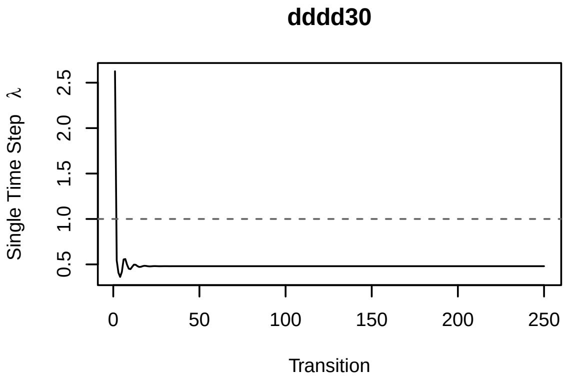
dddd30
Single Time Step λ
0.5 1.0 1.5 2.0 2.5
0 50 100 150 200 250
Transition

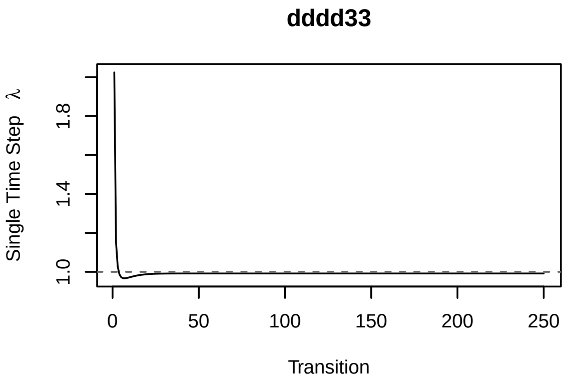
dddd33
Single Time Step λ
1.0 1.4 1.8
0 50 100 150 200 250
Transition

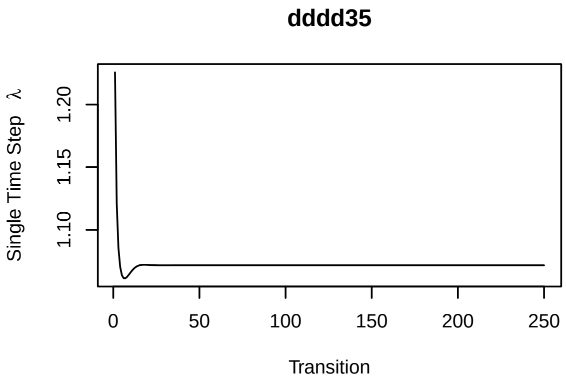
dddd35
Single Time Step λ
1.10 1.15 1.20
0 50 100 150 200 250
Transition

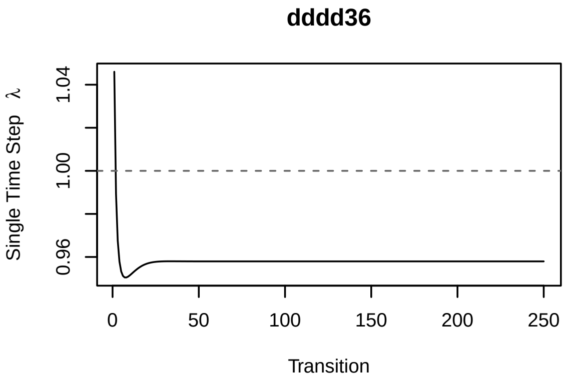
dddd36
Single Time Step λ
0.96 1.00 1.04
0 50 100 150 200 250
Transition

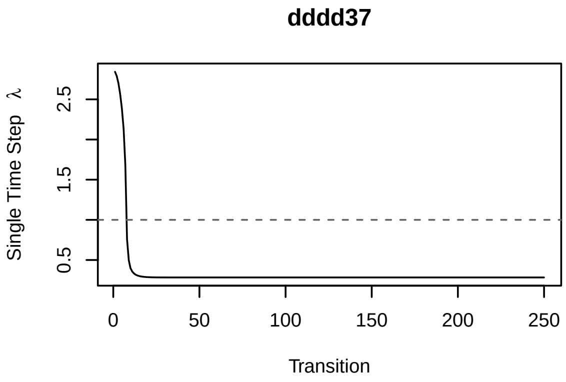
dddd37
Single Time Step λ
0.5 1.5 2.5
0 50 100 150 200 250
Transition

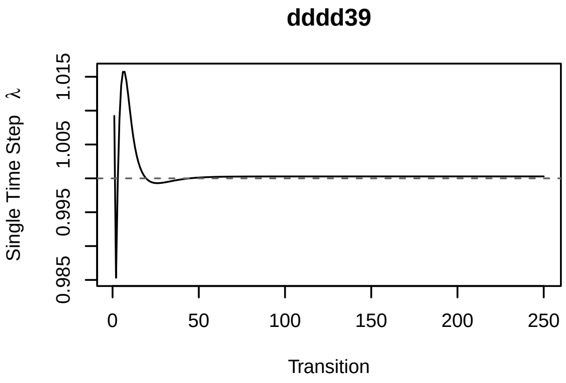
dddd39
Single Time Step λ
1.015
1.005
0.995
0.985
0
50
100
150
200
250
Transition

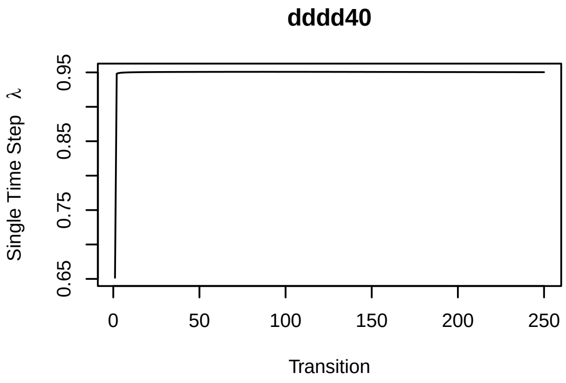
dddd40
Single Time Step λ
0.95
0.85
0.75
0.65
0
50
100
150
200
250
Transition

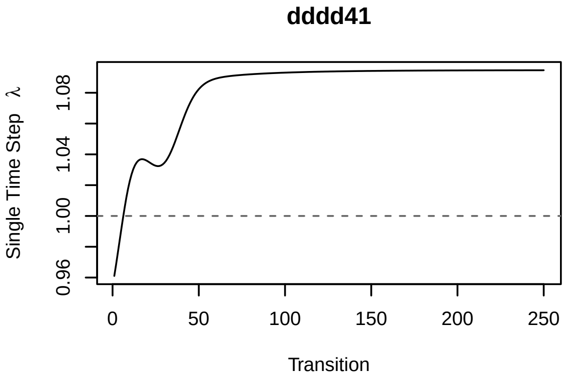
dddd41
Single Time Step λ
1.08
1.04
1.00
0.96
0
50
100
150
200
250
Transition

`dddd24`, `dddd26`, `dddd40` and `dddd41` are still not converging. We will remove those from our further analyses:

```
keep_ind <- setdiff(names(new_ipms), c("dddd24", "dddd26",
                                       "dddd40", "dddd41"))

new_ipms <- new_ipms[keep_ind]
```

### Further analyses

*Rpadrino* contains methods for most of *ipmr*'s analysis functions. These include (but are not limited to!) `lambda`, `left_ev`, and `right_ev`. We need all three of these to compute sensitivity and elasticity. We also need the binwidth of the integration mesh so we can perform integrations. *Rpadrino*'s has the `int_mesh()` function for that, and the binwidth is always the first element in the list that it returns. We can extract them like so:

```
lambdas    <- lambda(new_ipms)
repro_vals <- left_ev(new_ipms, tolerance = 1e-5)
ssd_vals   <- right_ev(new_ipms, tolerance = 1e-5)

d_zs <- lapply(new_ipms, function(x) int_mesh(x, full_mesh = FALSE)[[1]])
```

#### Sensitivity

With these, we can now compute sensitivity. This is given by $\mathbf{s}(z_0', z_0) = \frac{v(z_0')w(z_0)}{\langle v,w \rangle}$, where $v(z_0')$ is the left eigenvector and $w(z_0)$ is the right eigenvector. It will be helpful to write a function that takes these values as arguments and returns the sensitivity kernel. We will use `lapply(seq_along())` to iterate over each model.

```
# r_evs: right eigenvectors
# l_evs: left eigenvectors
# d_zs:  binwidths
# lapply(seq_along()) generates a sequence of numbers that correspond to indices
# in the list of eigenvectors and binwidths. Since each of these objects is a
# list of lists, we need to use [[index]][[1]]. The first[[]] gets the correct list
# entry, and the second [[1]] converts it to a numeric vector by unlisting the
# second layer of the list.

sens <- function(r_evs, l_evs, d_zs) {

  lapply(seq_along(r_evs),
         function(ind, r_ev, l_ev, d_z){

           outer(l_ev[[ind]][[1]], r_ev[[ind]][[1]]) /
             (sum(l_ev[[ind]][[1]] * r_ev[[ind]][[1]] * d_z[[ind]]))

         },
         r_ev = r_evs,
         l_ev = l_evs,
         d_z = d_zs)
}
```

```
sens_list <- sens(ssd_vals, repro_vals, d_zs) %>%
  setNames(names(lambdas))
```

We can plot these using `image()`:

```
par(mfrow = c(1, 1))
image(t(sens_list$aaa385))
```

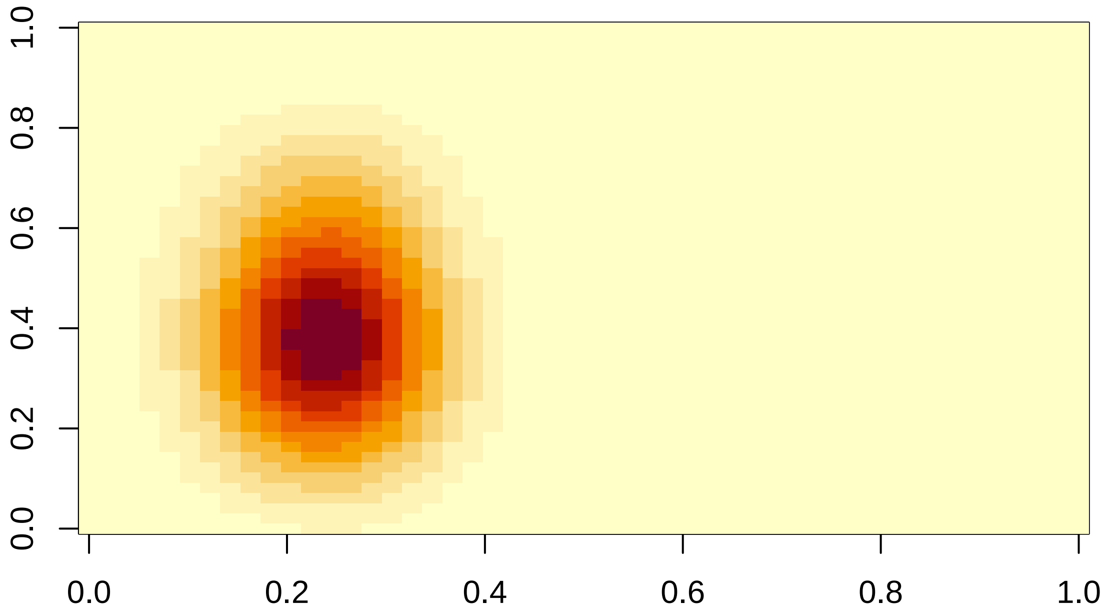

#### Elasticity

We can compute elasticity without much more effort. We need one more piece of information from the IPM list that we have not extracted - the iteration kernel. We can get those using `make_iter_kernel()` on the `new_ipms` object, and then computing the elasticity using $\mathbf{e}(z_0', z_0) = \frac{K(z_0', z_0)}{\lambda}\mathbf{s}(z_0', z_0)$.

```
iter_kerns <- make_iter_kernel(new_ipms)

elas_list <- lapply(seq_along(iter_kerns),
                    function(ind, iter_kernels, sens_kernels, lambdas, d_zs) {

                      (iter_kernels[[ind]][[1]] / d_zs[[ind]] / lambdas[[ind]]) *
                        sens_kernels[[ind]]

                    },
                    iter_kernels = iter_kerns,
                    sens_kernels = sens_list,
                    lambdas = lambdas,
```

```r
                    d_zs = d_zs) %>%
  setNames(names(lambdas))
```

Similarly, we can plot this using `image()`:

```r
image(t(elas_list$aaa385))
```

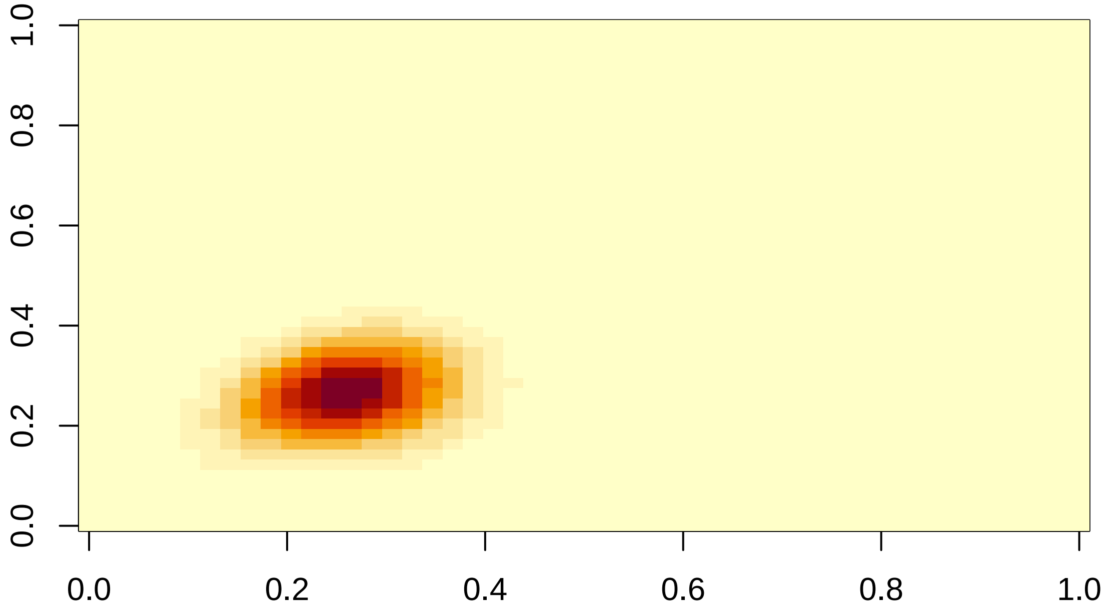

#### Mean lifetime recruit production

We can calculate the expected number of recruits produced over an individuals lifetime as function of its initial size. This is defined as $\bar{r}(z_0) = eFN$. $F$ is the fecundity kernel, and we can get these kernels from each IPM in our list using:

```r
F_kerns <- lapply(new_ipms, function(x) x$sub_kernels$F)
```

$N(z', z_0)$ is the fundamental operator. This tells us the expected amount of time an individual will spend in state $z'$ given an initial state $z_0$. The fundamental operator is defined as $(I - P)^{-1}$ (see Ellner, Childs, & Rees 2016 Chapter 3 for the derivation of this). $I$ is an identity kernel ($I(z', z) = 1$ for $z' = z$, and 0 everywhere else). This code is only a bit more complicated:

```r
# Function to create an identity kernel with dimension equal to P
make_i <- function(P) {
  return(
    diag(nrow(P))
  )
}
```

```r
N_kerns <- lapply(new_ipms, function(x) {

  P <- x$sub_kernels$P
  I <- make_i(P)

  # solve() inverts the matrix for us (the ^(-1) part of the equation)
  solve(I - P)

})
```

$e$ is a constant function $e(z) \equiv 1$. In practice, the left multiplication of $eF$ has the effect of computing the column sums of $F$. We will replace the $e$ with a call to `colSums()` in our code below (this will run faster than doing the multiplication). We now have everything we need to compute and visualize the expected lifetime reproductive output:

```r
# We wrap the computation in as.vector so that it returns a simple numeric vector
# rather than a 1 x N matrix

r_bars <- lapply(seq_along(N_kerns),
                 function(idx, Fs, Ns) {

                   as.vector(colSums(Fs[[idx]]) %*% Ns[[idx]])

                 },
                 Fs = F_kerns,
                 Ns = N_kerns) %>%
  setNames(names(F_kerns))

# we will extract the meshpoint values so that the x-axes on our plots look
# prettier.
x_seqs <- lapply(new_ipms,function(x) int_mesh(x, full_mesh = FALSE)[[2]])

# Finally, we can plot the data by looping over the lists and creating a
# a simple line plot (type = "l")

par(mfrow = c(4, 2))

for(i in seq_along(r_bars)) {

  plot(r_bars[[i]], x = x_seqs[[i]], type = 'l', main = names(r_bars)[i],
       ylab = expression(bar(r)(z[0])),
       xlab = expression(z[0]))

}
```

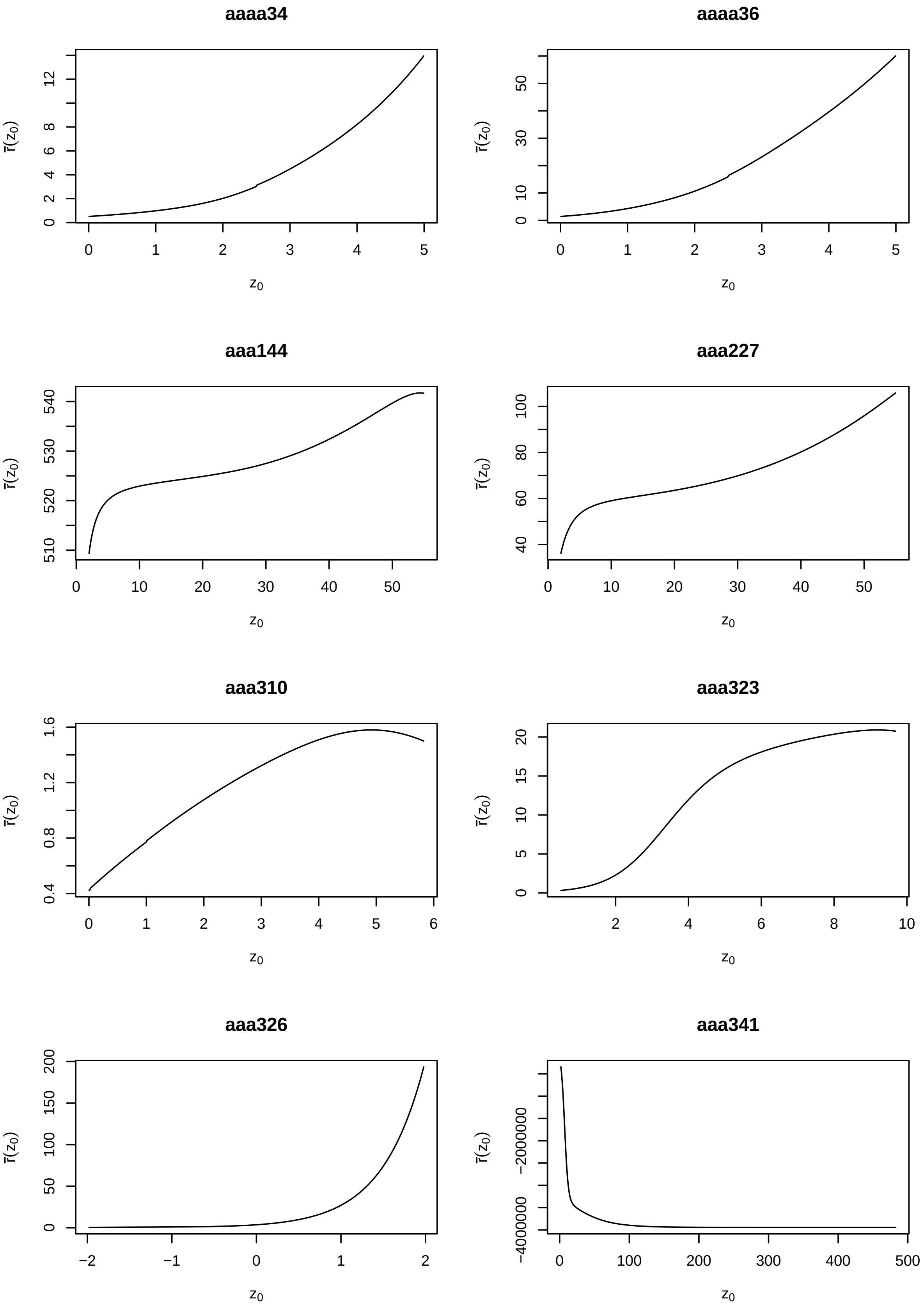

aaaa34
aaaa36
aaa144
aaa227
aaa310
aaa323
aaa326
aaa341
$\bar{r}(z_0)$
$z_0$

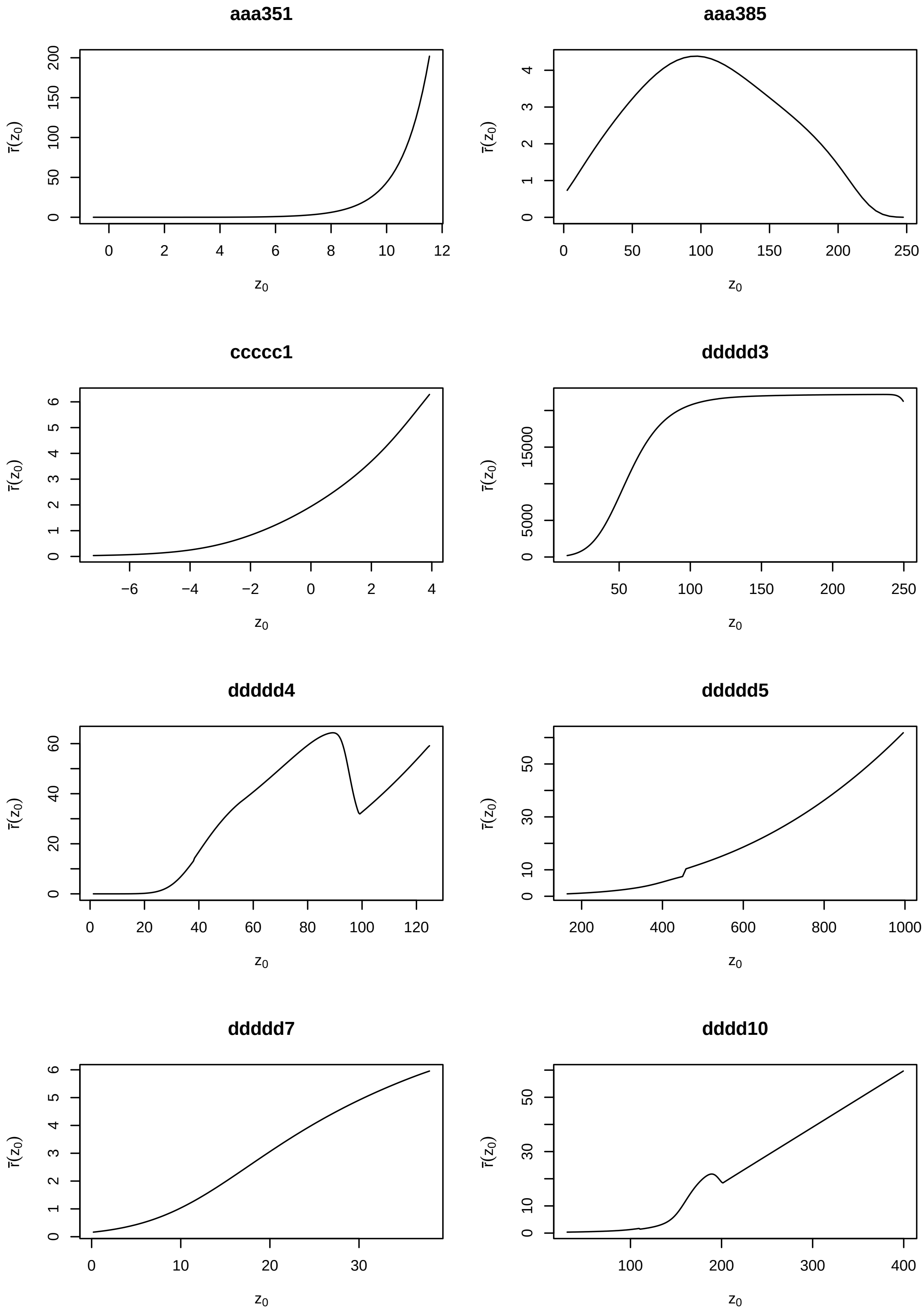
aaa351
aaa385
ccccc1
ddddd3
ddddd4
ddddd5
ddddd7
dddd10
$\bar{r}(z_0)$
$z_0$

**dddd29**

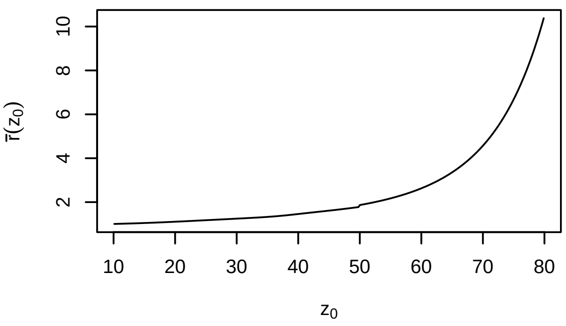

**dddd30**

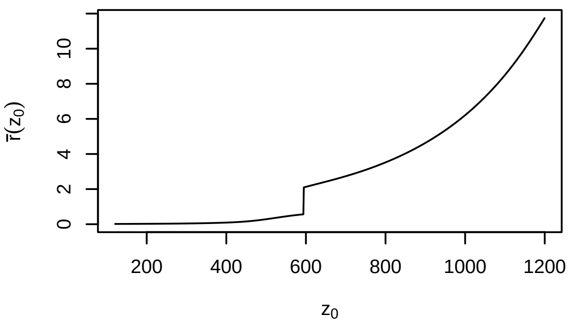

**dddd33**

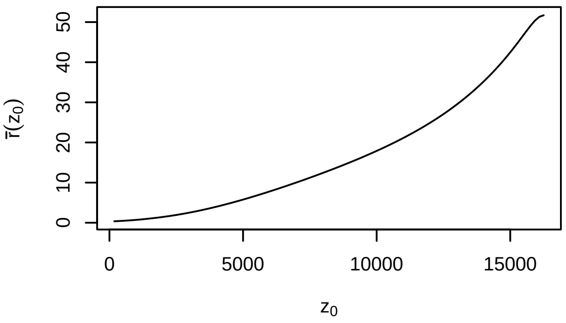

**dddd35**

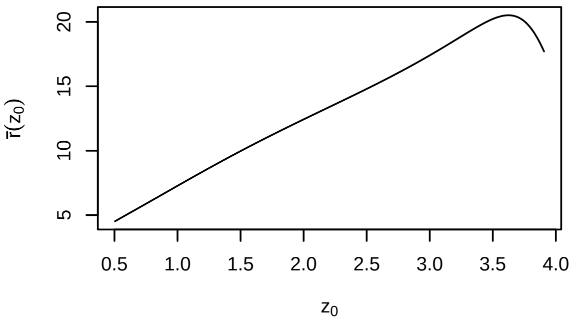

**dddd36**

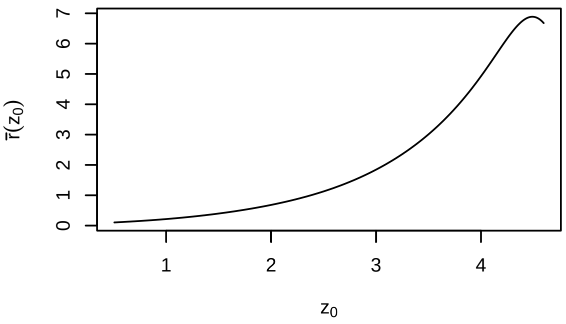

**dddd37**

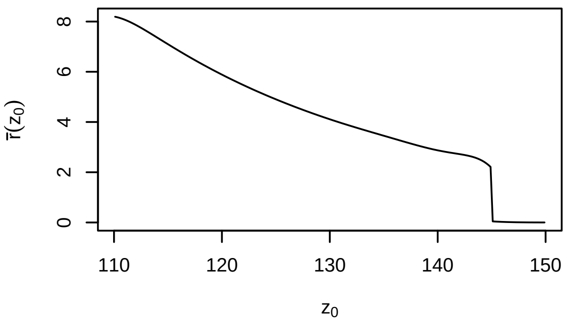

**dddd39**

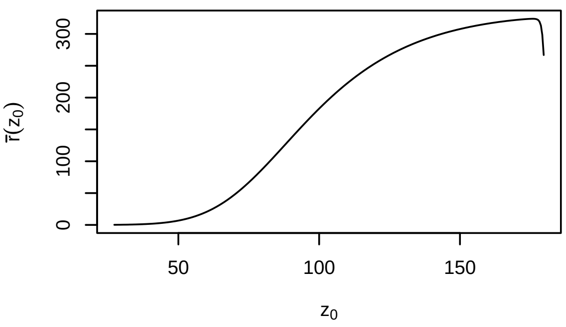

#### Troubleshooting and Modifying IPMs

The plot of the output above provides a cautionary tale: $\bar{r}(z_0)$ is negative in model aaa341! This is not biologically possible - we can not make negative recruits. We know there is some quirk in that model that produces these results. PADRINO would flag a model with negative numbers in the $F$ kernel when it's built, so we need to check the $N$ kernel for that model for negative values.

```
range(N_kerns$aaa341)
```

```
## [1] -5008.038563     1.020386
```

We have found the problem in the $N$ kernel, but what causes that? It turns out that the $P$ kernel, when discretized, is either close to or exactly singular, and so the determinant $(I - P)$ is very close to singular as well (and negative!). This can happen when the survival function is exactly 1 for some range of initial trait values (Ellner, Childs & Rees 2016, Chapter 3).

```
P <- new_ipms$aaa341$sub_kernels$P
I <- make_i(P)
det(I - P) # negative and very close to 0
```

```
## [1] -1.29379e-09
```

It is important to remember that PADRINO provides models *as they are published*, and does not try to correct these problems. Thus, data in here can cause problems if not treated with care!

We can try to modify the model very slightly to see if we can make this kernel non-singular. We will try to set a parallel minimum for the `s` function value in that model, which will hopefully pull our $det(I - P)$ into positive territory. We need to work with the `proto_ipm` object for this, because we want to propagate the function value changes to the sub-kernels (*i.e.* the P kernel).

We can alter the `s` function with `vital_rate_exprs<-` and `pdb_new_fun_form()`. We will update it to take the parallel minimum of the published `s` function, and the maximum value we'd want the function to ideally have (in this case, 0.98).

```
# First, peak at the current functional form
vital_rate_exprs(simple_det_list)$aaa341
```

```
## s: 1/(1 + exp(-(si + ss1 * size_1 + ss2 * size_1^2)))
## g_mean: gi + gs * size_1
## g: stats::dnorm(size_2, g_mean, g_sd)
## Fp: 1/(1 + exp(-(fpi + fps * size_1)))
## Fs: exp(fi + fs * size_1)
## Fd: stats::dnorm(size_2, fd_mean, fd_sd)
```

```
# Now, update it with our desired maximum value
vital_rate_exprs(simple_det_list) <- pdb_new_fun_form(
  list(
    aaa341 = list(
      s = pmin(0.98, 1/(1 + exp(-(si + ss1 * size_1 + ss2 * size_1^2))))
    )
  )
)

# we will skip rebuilding the whole list - we just want to make sure we have fixed
# this particular model. The species in this model is Lonicera maackii.

new_lonicera <- pdb_make_ipm(simple_det_list["aaa341"])
```

```r
P <- new_lonicera$aaa341$sub_kernels$P
I <- make_i(P)

cat("New det(I - P) is: ", det(I - P))
```

```
## New det(I - P) is:  1.22853e-05
```

```r
N <- solve(I - P)
F <- new_lonicera$aaa341$sub_kernels$F

r_bar_lonicera <- as.vector(colSums(F) %*% N)

plot(r_bar_lonicera,
     type = 'l',
     ylab = expression(bar(r)(z[0])),
     xlab = expression(z[0]))
```

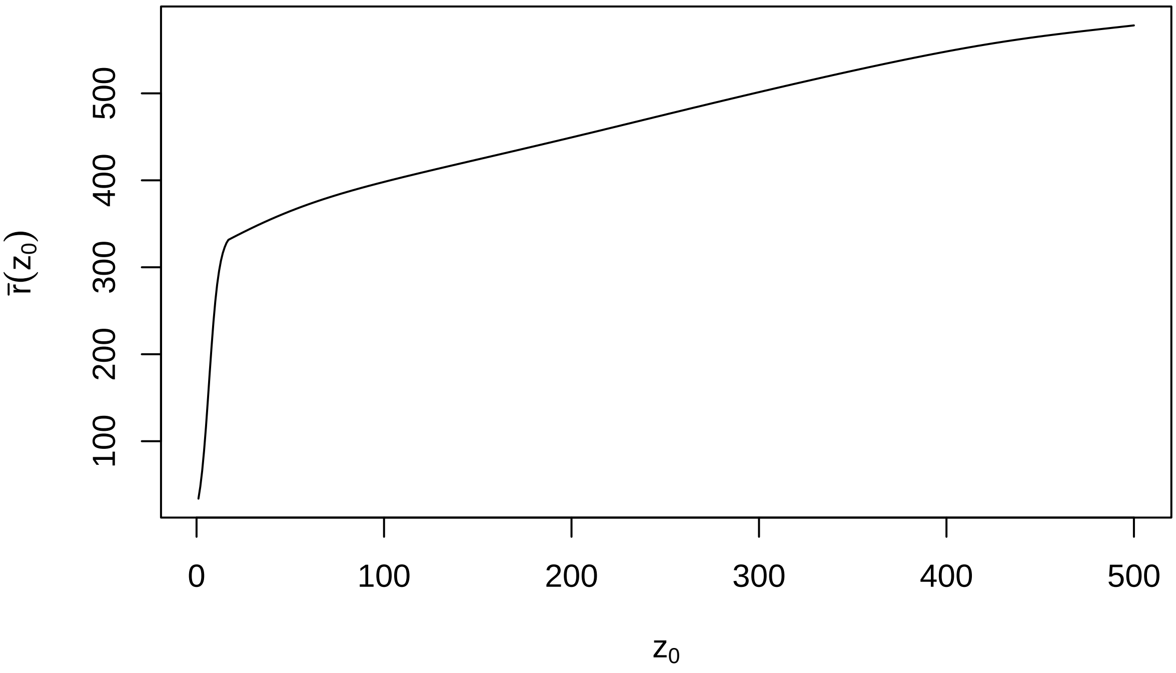

```r
# Since we are now happier with how this IPM is behaving, we will insert it into
# our list for subsequent analyses
new_ipms$aaa341 <- new_lonicera$aaa341
```

That looks a bit closer to reality!

### Vital rate and parameter level analyses

We can also run analyses at the parameter and the vital rate level for PADRINO. These require more care - it is strongly recommended to check the original publications to for the meaning of each parameter and vital rate. There is simply too much variability in the way vital rates and parameters are estimated in the literature to provide systematic descriptions of them in PADRINO. With this caveat in mind, we will proceed to a couple examples of using vital rate functions and parameters in further analyses.

The ability to perturb function values and parameter estimates is one of the great strengths of IPMs. Furthermore, computing many life history traits requires the values of vital rate functions. Therefore, we took great pains when designing the database to ensure these analyses were still possible. As noted above, they require some additional effort, but are usually worth it. We will step through the code pieces required to extract these below. We will start with an example computing the sensitivity of $\lambda$ to vital rate function values. After that, we will show how to compute the mean size at death conditional on initial state $z_0$ and the size at death kernel $\Omega(z', z_0)$, both of which rely on extracting the survival functions.

#### Vital rate function value perturbations

These require modifying the general sensitivity formula to compute the partial derivative of $\lambda$ with respect to change in $f(z)$. These expressions depend on the form of the kernel, and so no general formula exists for function value perturbations. However, we can use the chain rule and the general formula for sensitivity to a given perturbation to work it out. The latter formula is:

1. $\frac{\partial\lambda}{\partial\epsilon}\bigg|_{\epsilon=0} = \frac{\langle v, Cw \rangle}{\langle v, w \rangle}$.

Here, $v$ and $w$ are the left and right eigenvectors of the iteration kernel (provided by `left_ev` and `right_ev`), and $C$ is the perturbation kernel, which we will need to identify. We will take the following steps:

1. Identify models we want to use.
2. Inspect the kernel formulae and vital rate functions using `print` methods.
3. Write down the perturbation kernels for each model.
4. Construct the IPM objects and extract $v$ and $w$.
5. Implement the perturbations in *R*.

##### Identifying models (1)

In order to keep things simple, we will work with models where survival only occurs in 1 kernel. There are numerous examples of how to extend these analyses elsewhere (*e.g.* Ellner, Childs & Rees 2016). We can look into this using the `pdb$VitalRateExpr$kernel_id` column in conjunction with the `pdb$VitalRateExpr$demographic_paratemer` column.

```
# Find all rows in VitalRateExpr corresponding to survival
init_ind <-
  my_pdb$VitalRateExpr[
    my_pdb$VitalRateExpr$demographic_parameter == "Survival", ]

# Next, select ipm_id's that have survival functions that only show up
# in "P"

keep_ind <- init_ind$ipm_id[init_ind$kernel_id == "P"] %>%
  unique()
```

```r
# To keep things quick, we will just use the first 3 in this index

keep_ind <- keep_ind[1:3]

vr_sens_pdb <- pdb_subset(my_pdb, keep_ind)
```

**Inspect the kernels and vital rate expressions (2)**

Next, we will construct `proto_ipm` objects for each model and check to see how the kernels are constructed:

```r
proto_list <- pdb_make_proto_ipm(vr_sens_pdb)

proto_list[[1]]
```

```
## A simple, density independent, deterministic proto_ipm with 2 kernels defined:
##  P, F
##
## Kernel formulae:
##
## P: s * g
## F: r * fn * pE * d
##
## Vital rates:
##
## s: 1/(1 + exp(-(s_b + s_m * lnsize_1)))
## g_mean: g_b + g_m * lnsize_1
## g_var: sqrt(gv_b + gv_m * lnsize_1)
## g: stats::dnorm(lnsize_2, g_mean, g_var)
## r: 1/(1 + exp(-(r_b + r_m * lnsize_1)))
## fn: exp(fn_b + fn_m * lnsize_1)
## d: stats::dexp(lnsize_2, 1/d_mean)
##
## Parameter names:
##
##  [1] "s_b"    "s_m"    "g_b"    "g_m"    "gv_b"   "gv_m"   "r_b"    "r_m"
##  [9] "fn_b"   "fn_m"   "t_r"    "d_mean" "pE"
##
## All parameters in vital rate expressions found in 'data_list':  TRUE
##
## Domains for state variables:
##
## lnsize: lower_bound = 0, upper_bound = 5, n_meshpoints = 500
##
## Population states defined:
##
## n_lnsize: Pre-defined population state.
##
## Internally generated model iteration procedure:
##
## n_lnsize_t_1: right_mult(kernel = P, vectr = n_lnsize_t) + right_mult(kernel = F,
##     vectr = n_lnsize_t)
```

```r
proto_list[[2]]
```

```
## A simple, density independent, deterministic proto_ipm with 2 kernels defined:
##  P, F
##
## Kernel formulae:
##
## P: s * g
## F: r * fn * pE * d
##
## Vital rates:
##
## s: 1/(1 + exp(-(s_b + s_m * lnsize_1)))
## g_mean: g_b + g_m * lnsize_1
## g_var: sqrt(gv_b + gv_m * lnsize_1)
## g: stats::dnorm(lnsize_2, g_mean, g_var)
## r: 1/(1 + exp(-(r_b + r_m * lnsize_1)))
## fn: exp(fn_b + fn_m * lnsize_1)
## d: stats::dexp(lnsize_2, 1/d_mean)
##
## Parameter names:
##
##  [1] "s_b"    "s_m"    "g_b"    "g_m"    "gv_b"   "gv_m"   "r_b"    "r_m"
##  [9] "fn_b"   "fn_m"   "t_r"    "d_mean" "pE"
##
## All parameters in vital rate expressions found in 'data_list':  TRUE
##
## Domains for state variables:
##
## lnsize: lower_bound = 0, upper_bound = 5, n_meshpoints = 500
##
## Population states defined:
##
## n_lnsize: Pre-defined population state.
##
## Internally generated model iteration procedure:
##
## n_lnsize_t_1: right_mult(kernel = P, vectr = n_lnsize_t) + right_mult(kernel = F,
##     vectr = n_lnsize_t)
```

```
proto_list[[3]]
```

```
## A simple, density independent, deterministic proto_ipm with 2 kernels defined:
##  P, F
##
## Kernel formulae:
##
## P: s * g
## F: rep_p * es_p * sdl_s * n_infl * n_fl * n_seed
##
## Vital rates:
##
## s: ssurv * wsurv
## ssurv: exp(ssurv_i + ssurv_s * ((log(size_1^3) - smlv)/sslv) + ssurv_el *
##     elev_s)/(1 + exp(ssurv_i + ssurv_s * ((log(size_1^3) - smlv)/sslv) +
##     ssurv_el * elev_s))
## wsurv: exp(wsurv_i + wsurv_s * ((log(size_1^3) - wmlv)/wslv) + wsurv_f *
```

```
##     cumfrost)/(1 + exp(wsurv_i + wsurv_s * ((log(size_1^3) -
##     wmlv)/wslv) + wsurv_f * cumfrost))
## g_mean: sign(growth) * abs(growth)^(1/3)
## growth: g_i + g_el * elev_g + g_frost * annfrost + g_s * (size_1^3)
## g: stats::dnorm(size_2, g_mean, g_sd)
## rep_p: fl_p * germ_p
## fl_p: exp(fl_i + fl_s * ((log(size_1^3) - fmlv)/fslv))/(1 + exp(fl_i +
##     fl_s * ((log(size_1^3) - fmlv)/fslv)))
## germ_p: exp(germ_i + germ_el * elev_germ)/(1 + exp(germ_i + germ_el *
##     elev_germ))
## n_infl: exp(infl_n)
## n_fl: exp(fl_n)
## n_seed: exp(seed_i)
## sdl_s: stats::dnorm(size_2, sdl_mean, sdl_sd)
##
## Parameter names:
##
##  [1] "ssurv_i"   "ssurv_el"  "ssurv_s"   "wsurv_i"   "wsurv_s"   "wsurv_f"
##  [7] "g_i"       "g_el"      "g_frost"   "g_s"       "g_sd"      "fl_i"
## [13] "fl_s"      "germ_i"    "germ_el"   "es_p"      "sdl_mean"  "sdl_sd"
## [19] "infl_n"    "fl_n"      "seed_i"    "smlv"      "sslv"      "wmlv"
## [25] "wslv"      "fmlv"      "fslv"      "annfrost"  "cumfrost"  "elev_germ"
## [31] "elev_g"    "elev_s"
##
## All parameters in vital rate expressions found in 'data_list':  TRUE
##
## Domains for state variables:
##
## size: lower_bound = 2, upper_bound = 55, n_meshpoints = 1000
##
## Population states defined:
##
## n_size: Pre-defined population state.
##
## Internally generated model iteration procedure:
##
## n_size_t_1: right_mult(kernel = P, vectr = n_size_t) + right_mult(kernel = F,
##     vectr = n_size_t)
```

We can see that for each IPM, $P = s(z) * G(z', z)$. In the third models, $s(z)$ is comprised of two additional functions, which we can perturb individually. We will show a quick example of that after applying our perturbation kernels to all 3.

##### Write out the perturbation kernels (3)

Our perturbation kernel for all 3 models will take the following form: $C(z', z) = \delta_{z_0}(z)G(z', z)$. With a little rearranging (see Ellner, Childs, & Rees 2016 Chapter 4), we find the following:

$$\frac{\partial\lambda}{\partial s(z_0)} = \frac{\int v(z')G(z', z_0)w(z_0)dz'}{\int v(z)w(z)dz} = \frac{(vG) \circ w}{\langle v, w\rangle}.$$

The second portion of the equations above is the first part re-written to use operator notation (which drops the $z$s and $z'$s for brevity). The $\circ$ denotes point-wise multiplication.

##### Implement the models (4)

Now that we have written down our perturbation formulae, we need to rebuild the models, and make use of some non-standard arguments to `pdb_make_ipm`. By default, *Rpadrino* does not return the vital rate functions values. To get those, we need to specify `return_all_envs = TRUE` in the `addl_args` list.

```
arg_list <- lapply(keep_ind, function(x) list(return_all_envs = TRUE)) %>%
  setNames(keep_ind)

ipm_list <- pdb_make_ipm(proto_list, addl_args = arg_list)

r_evs    <- right_ev(ipm_list)
l_evs    <- left_ev(ipm_list)
```

##### Implement the perturbations (5)

Next, we need to implement the formula above. This is fairly straightforward, and we will make use of another function in *Rpadrino*: `vital_rate_funs()` (not to be confused with `vital_rate_exprs()`!). This extracts the vital rate function values from each model and returns them in a named list (`vital_rate_exprs()` extracts the expressions that create these values). Let's see what this looks like:

```
vr_funs <- vital_rate_funs(ipm_list)

vr_funs$aaaa34
```

```
## $P
## s (not yet discretized): A 500 x 500 kernel with minimum value: 0.3638 and maximum value: 0.852
## g_mean (not yet discretized): A 500 x 500 kernel with minimum value: 1.1114 and maximum value: 3.6806
## g_var (not yet discretized): A 500 x 500 kernel with minimum value: 0.9749 and maximum value: 0.9749
## g (not yet discretized): A 500 x 500 kernel with minimum value: 0 and maximum value: 0.1293
##
## $F
## r (not yet discretized): A 500 x 500 kernel with minimum value: 0.0062 and maximum value: 0.9969
## fn (not yet discretized): A 500 x 500 kernel with minimum value: 43.05 and maximum value: 1280.9888
## d (not yet discretized): A 500 x 500 kernel with minimum value: 0 and maximum value: 0.1481
```

```
vr_funs$aaa144
```

```
## $P
## s (not yet discretized): A 1000 x 1000 kernel with minimum value: 0.9712 and maximum value: 0.9997
## ssurv (not yet discretized): A 1000 x 1000 kernel with minimum value: 0.9729 and maximum value: 0.999
## wsurv (not yet discretized): A 1000 x 1000 kernel with minimum value: 0.9983 and maximum value: 1
## g_mean (not yet discretized): A 1000 x 1000 kernel with minimum value: 18.6951 and maximum value: 42
## growth (not yet discretized): A 1000 x 1000 kernel with minimum value: 6534.029 and maximum value: 75
## g (not yet discretized): A 1000 x 1000 kernel with minimum value: 0 and maximum value: 0.0798
##
## $F
## rep_p (not yet discretized): A 1000 x 1000 kernel with minimum value: 0 and maximum value: 0.0049
## fl_p (not yet discretized): A 1000 x 1000 kernel with minimum value: 0 and maximum value: 0.3948
## germ_p (not yet discretized): A 1000 x 1000 kernel with minimum value: 0.0124 and maximum value: 0.01
## n_infl (not yet discretized): A 1000 x 1000 kernel with minimum value: 2.4843 and maximum value: 2.48
## n_fl (not yet discretized): A 1000 x 1000 kernel with minimum value: 131.6307 and maximum value: 131
## n_seed (not yet discretized): A 1000 x 1000 kernel with minimum value: 13.1971 and maximum value: 13
## sdl_s (not yet discretized): A 1000 x 1000 kernel with minimum value: 0 and maximum value: 0.3988
```

We see from the printed values that each vital rate function contains the complete $n \times n$ set of values for each combination of meshpoints. Additionally, it warns us that these are not yet integrated. This is actually a good thing - we want the continuous function values for the sensitivity, not the discretized values. We know from our formula above that we need to extract $G(z', z)$, and that these are named `g` in each model. This will not always be true for PADRINO, so care must be taken at this step to make sure you extract the correct values!

Recall that we are about to implement: $\frac{(vG) \circ w}{\langle v,w \rangle}$. The $\langle ... \rangle$ is the inner product of $v, w$, and so we also need the value of $dz$ to implement the denominator. We will get those using `int_mesh()` again.

```
mesh <- lapply(ipm_list, function(x) int_mesh(x, full_mesh = FALSE))
d_zs <- lapply(mesh, function(x) x[[1]])

sens_list <- lapply(seq_along(vr_funs),
                    function(idx, r_evs, vr_funs, l_evs, d_zs) {

                      # Extract objects to temporary values so the formula is more
                      # readable

                      G <- vr_funs[[idx]]$P$g
                      v <- unlist(l_evs[[idx]])
                      w <- unlist(r_evs[[idx]])
                      d_z <- d_zs[[idx]]

                      numerator <- as.vector((v %*% G) * w)
                      denominator <- sum(v * w * d_z)

                      numerator / denominator

                    },
                    r_evs = r_evs,
                    l_evs = l_evs,
                    vr_funs = vr_funs,
                    d_zs = d_zs)

par(mfrow = c(3, 1))

for(i in seq_along(sens_list)) {

  plot(sens_list[[i]], x = mesh[[i]][[2]],
       type = "l",
       main = names(mesh)[i],
       ylab = expression(bold(s)(z[0])),
       xlab = expression(z[0]))

}
```

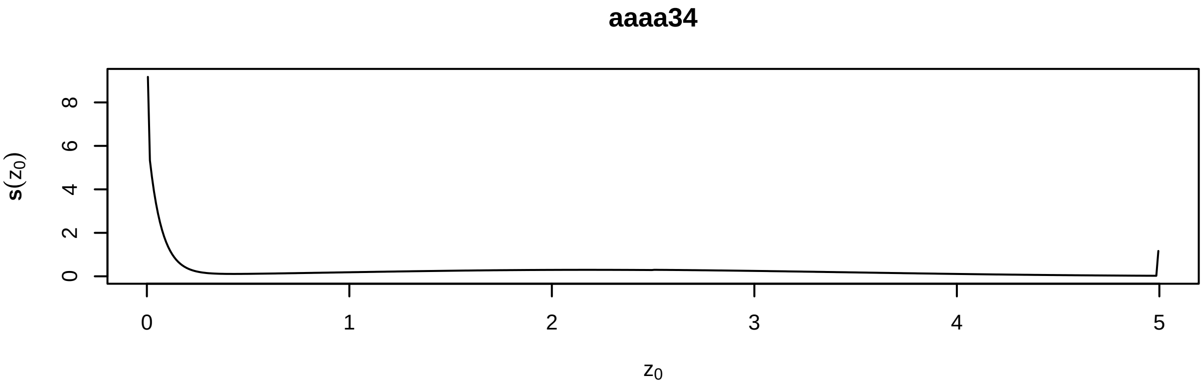

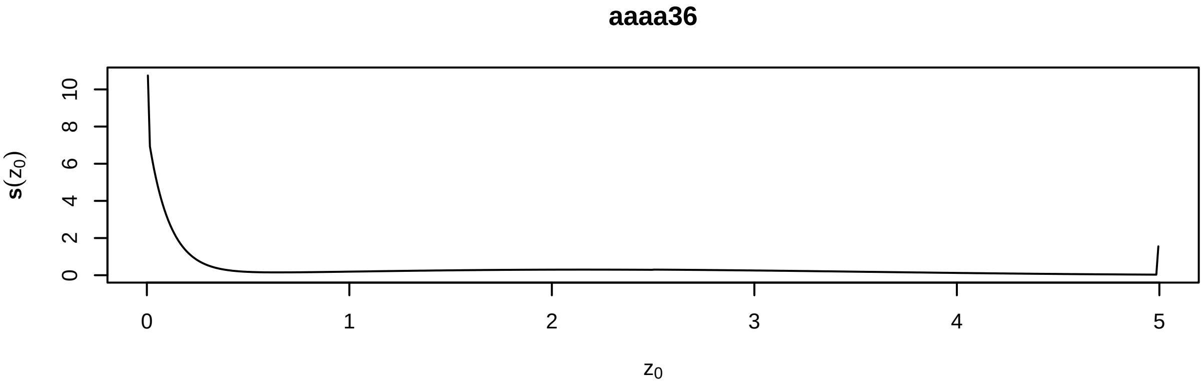

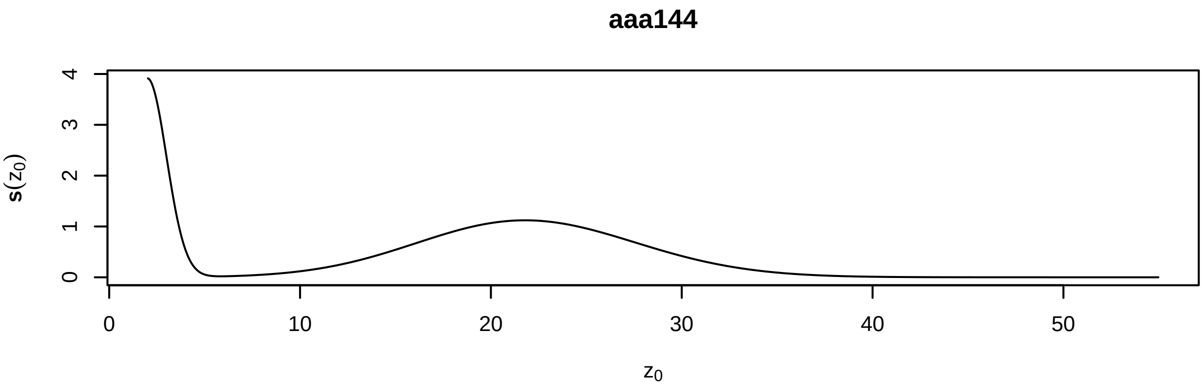

##### Other perturbation kernels

As mentioned above, the survival function in the last IPM is comprised of two additional functions: `ssurv` and `wsurv`. Thus, we could re-write the $P$ kernel as $P(z', z) = s_s(z) {*} s_w(z) {*} G(z', z)$. Thus, if we wanted to know the effect of perturbing only $s_w(z)$, we would re-write our perturbation kernel as $C(z', z) = \delta(z_0) * s_s(z) * G(z', z)$, and our perturbation formula (in operator notation) becomes $\mathbf{s}_s(z_0) = \frac{(v s_s G) \circ w}{\langle v, w \rangle}$. We will drop the first model from our lists because this analysis does not apply to it. We can implement this by slightly modifying the code above:

```
s_s <- vr_funs[[3]]$P$ssurv
G <- vr_funs[[3]]$P$g
v <- unlist(l_evs[[3]])
w <- unlist(r_evs[[3]])
d_z <- d_zs[[3]]

numerator   <- as.vector((v %*% (s_s * G)) * w)
denominator <- sum(v * w * d_z)

sens_s_w <- numerator / denominator

par(mfrow = c(1, 1))

mesh_ps <- mesh[[3]][[2]]
nm      <- names(mesh)[3]

plot(y = sens_s_w, x = mesh_ps,
     type = "l",
     main = nm,
     ylab = expression(bold(s)[s[w]](z[0])),
     xlab = expression(z[0]))
```

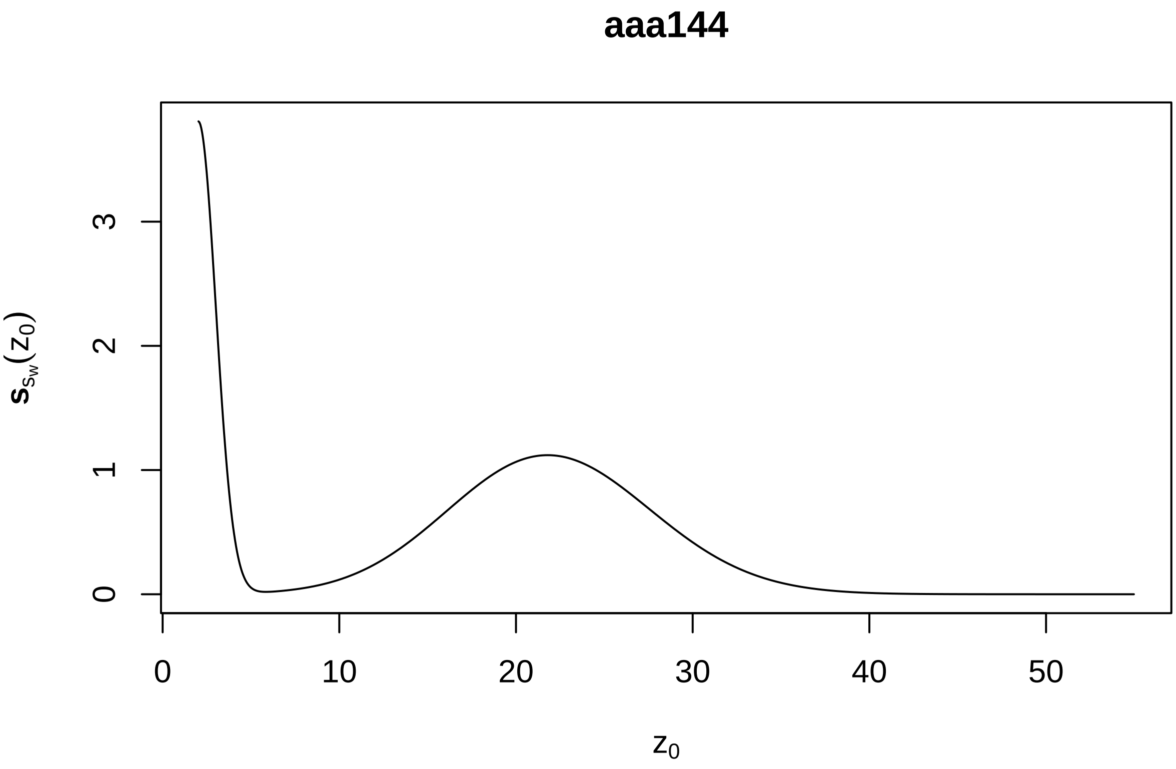

#### Mean size at death and size at death kernels

We can also use vital rate function values in conjunction with sub-kernels to implement calculations of life history traits. For this example, we will examine mean size at death and the size at death kernel. These are given by the following equations:

Mean size at death $= \bar{\omega} = (\mathbf{i} \circ (1 - s))N$ and size at death kernel $= \Omega(z', z_0) = (1 - s(z')) * N(z', z_0)$.

In these equations, $s$ and $s(z')$ are survival functions, and $N$ is the fundamental operator, which is defined as $N = (I - P)^{-1}$, where $P$ is the survival/growth kernel from the IPM and $I$ is an identity kernel (analogous to an identity matrix). Fortunately, we can reuse our `N_kerns` code from above. However, it is important to remember that the $(1 - s)$ term represents all mortality pathways, and species may have more than one way to die (*e.g.* monocarpic perennials die through natural mortality as well as the flowering process). Therefore, we also want to check our kernel formulae and see if those include additional terms that may represent alternative mortality pathways. Additionally, we need to get the meshpoints which correspond to $\mathbf{i}$.

This time, we will use considerably more IPMs - we will get those from the `new_ipms` object we created earlier. However, we need to rebuild them with `return_all_envs = TRUE` so that we can access the vital rate function values.

```
arg_list <- lapply(names(new_ipms), function(x) list(return_all_envs = TRUE,
                                                     iterations      = 250)) %>%
  setNames(names(new_ipms))

new_ipms <- pdb_make_ipm(simple_det_list, addl_args = arg_list)

# Removing our these problem IPMs we identified before

keep_ind <- setdiff(names(new_ipms), c("dddd24", "dddd26",
                                       "dddd40", "dddd41"))

new_ipms <- new_ipms[keep_ind]

kernel_formulae(new_ipms)
```

```
## $aaaa34
## P: s * g
## F: r * fn * pE * d
## $aaaa36
## P: s * g
## F: r * fn * pE * d
## $aaa144
## P: s * g
## F: rep_p * es_p * sdl_s * n_infl * n_fl * n_seed
## $aaa227
## P: s * g
## F: rep_p * es_p * sdl_s * n_infl * n_fl * n_seed
## $aaa310
## P: s * g
## F: f_n * f_d
## $aaa323
## P: s * g
## F: p_es * sdl_size * n_seeds
## $aaa326
## P: (1 - p_fl) * s * g
```

```
## F: p_fl * fec1 * sdl_size * est_p
## $aaa341
## P: s * g
## F: Ep * Fp * Fs * Fd
## $aaa351
## P: s * g
## F: Pf * Nfruit * Nseeds * Pe * Fd
## $aaa385
## P: s * g
## F: f * fd
## $ccccc1
## P: s * g
## F: p_r * r_s * r_d * r_r
## $ddddd3
## P: s * g
## F: d * r
## $ddddd4
## P: s * g
## F: d * r
## $ddddd5
## P: s * g
## F: d * r
## $ddddd7
## P: s * g
## F: d * r
## $dddd10
## P: s * g * d_len
## F: d * r
## $dddd29
## P: s * g
## F: (s * r * pHS * d)/2
## $dddd30
## P: s * g
## F: s * r * pg * 0.5 * d
## $dddd33
## P: s * g
## F: rp * f * EP * d
## $dddd35
## P: s * g
## F: p_fertile * polyps * fec * p_est * d
## $dddd36
## P: s * g
## F: p_fertile * polyps * fec * p_est * d
## $dddd37
## P: s * g
## F: d * r
## $dddd39
## P: s * g
## F: f1 * f2 * f3 * f4 * f5 * d * 0.5
```

Upon further inspection, model ID `aaa326` contains a `(1 p_fl)` term, which represents mortality due to flowering. We need to include this in the calculations above. We do this like so:

$\bar{\omega} = (\mathbf{i} * (p_{fl} + (1 - p_{fl}) * (1 - s))N,$

and

$\Omega(z', z_0) = (p_{fl}(z') + (1 - p_{fl}(z')) * (1 - s(z')))N(z', z_0)$.

This can be summarized by saying "a plant dies with flowering probability $p_{fl}$, and if the plant does not flower $(1 - p_{fl})$, it dies with probability $1 - s$." We are now ready to proceed with our calculations!

```
N_kerns <- lapply(new_ipms, function(x) {

  P <- x$sub_kernels$P
  I <- make_i(P)

  solve(I - P)

})

surv_funs <- lapply(new_ipms, function(x) {
  vital_rate_funs(x)$P$s
})

p_flower <- vital_rate_funs(new_ipms$aaa326)$P$p_fl
```

The **i** corresponds to the sizes in each IPM. Thus, we want to get the meshpoints, which we do with `int_mesh` like before. The `lapply(x[[2]])` extracts the value of `z_1`, as this is always the second entry in the list of 3 returned by `int_mesh` (and we do not need the `d_z` or `z_2` values).

```
i_vals <- int_mesh(new_ipms, full_mesh = FALSE) %>%
  lapply(function(x) x[[2]])
```

Now, we just have to implement the calculations:

```
omega_bar_z <- lapply(seq_along(new_ipms),
                      function(index, i_vals, surv_funs, N_kerns, p_flower) {

                        id <- names(i_vals)[index]
                        i <- i_vals[[index]]

                        # The survival function is represented as a bivariate
                        # function in ipmr even though it is actually a univariate
                        # function of z. Thus, we need to pull out the
                        # univariate form of it to ensure we get the correct
                        # result from (1 - s) (_i.e._ a vector, not an array!).
                        # The correct univariate form is given by the rows,
                        # as every column contains the same values.

                        s <- surv_funs[[index]][1, ]
                        N <- N_kerns[[index]]

                        if(id == "aaa326") {
                          # Same indexing as the survival function
                          p_fl <- p_flower[1, ]

                          out <- (i * (p_fl + (1-p_fl) * (1 - s))) %*% N
                        } else {

                          out <- (i * (1 - s)) %*% N
```

```r
                        }

                        return(out)
                      },
                      i_vals     = i_vals,
                      surv_funs = surv_funs,
                      N_kerns    = N_kerns,
                      p_flower   = p_flower) %>%
  setNames(names(i_vals))

Omega_z0_z <- lapply(seq_along(new_ipms),
                     function(index, surv_funs, N_kerns, p_flower) {
                       id <- names(surv_funs)[index]

                       s <- surv_funs[[index]][1, ]
                       N <- N_kerns[[index]]

                       if(id == "aaa326") {
                         p_fl <- p_flower
                         out <- (p_fl + (1-p_fl) * (1 - s)) %*% N
                       } else {

                         out <- (1 - s) * N
                       }

                       return(out)
                     },
                     surv_funs = surv_funs,
                     N_kerns    = N_kerns,
                     p_flower   = p_flower) %>%
  setNames(names(i_vals))

par(mfrow = c(4, 2))

for(i in seq_along(omega_bar_z)) {

  plot(x    = i_vals[[i]],
       y    = as.vector(omega_bar_z[[i]]),
       type = "l",
       main = names(i_vals)[i],
       xlab = expression(paste(z[0])),
       ylab = expression(paste(bar(omega),"(z)")))
}
```

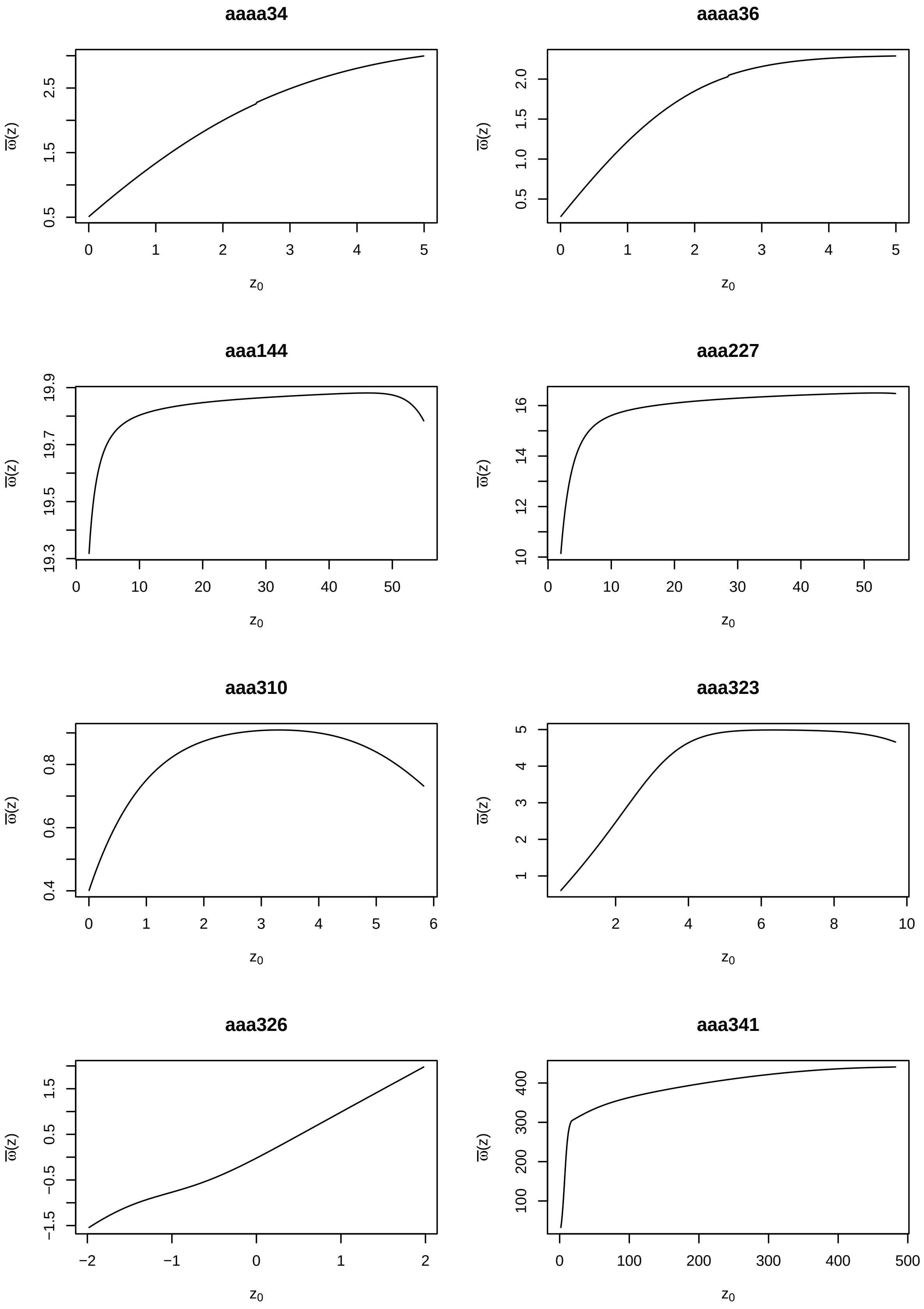
aaaa34
aaaa36
aaa144
aaa227
aaa310
aaa323
aaa326
aaa341
ϖ(z)
z0
0.5
1.5
2.5
0 1 2 3 4 5
0.5
1.0
1.5
2.0
19.3
19.5
19.7
19.9
0 10 20 30 40 50
10
12
14
16
0.4
0.6
0.8
0 1 2 3 4 5 6
1 2 3 4 5
2 4 6 8 10
−1.5
−0.5
0.5
1.5
−2 −1 0 1 2
100 200 300 400
0 100 200 300 400 500

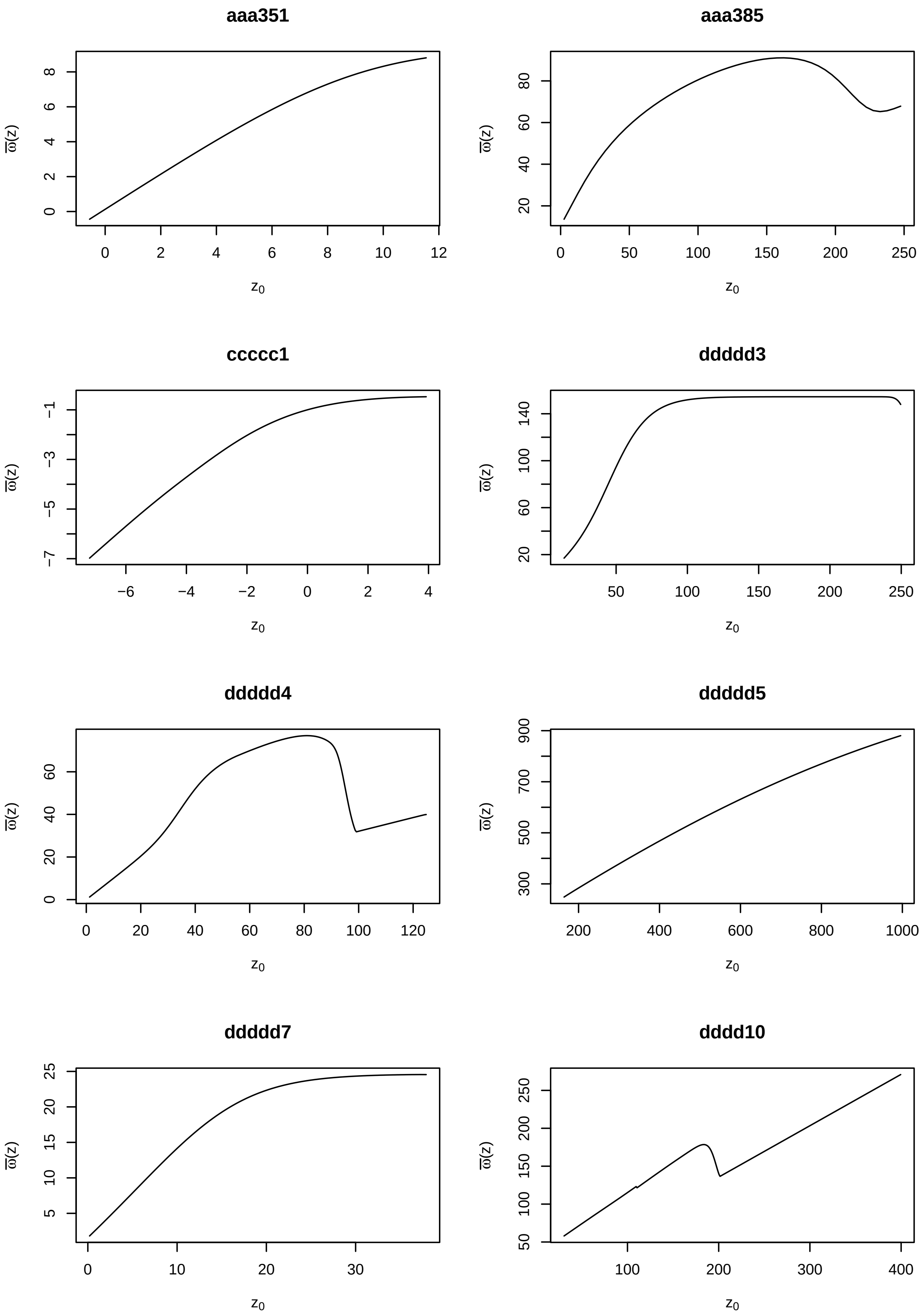
aaa351
aaa385
ccccc1
ddddd3
ddddd4
ddddd5
ddddd7
dddd10
$\overline{\varpi}(z)$
$z_0$

**dddd29**

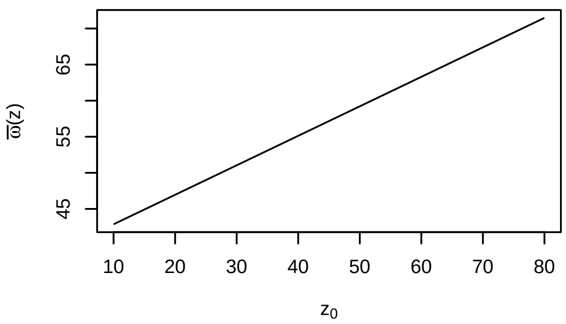

**dddd30**

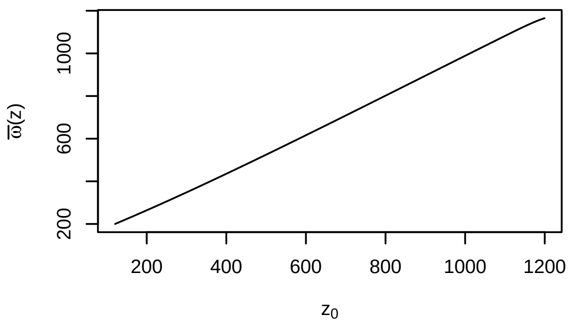

**dddd33**

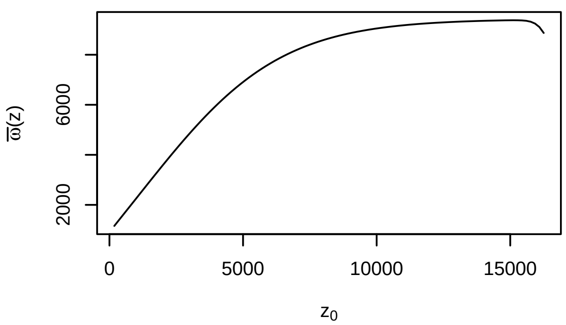

**dddd35**

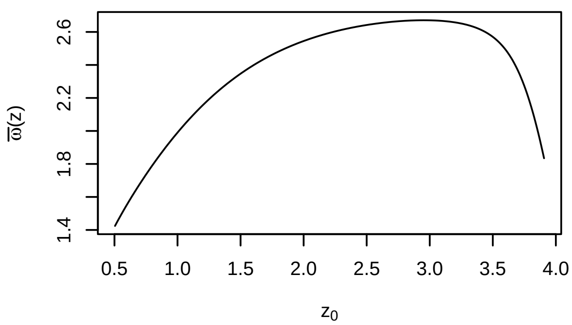

**dddd36**

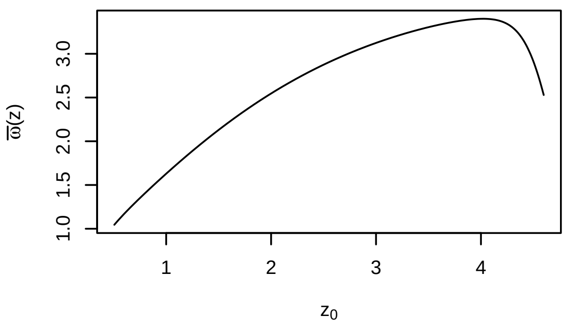

**dddd37**

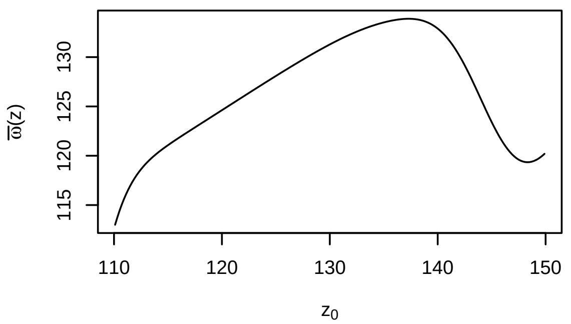

**dddd39**

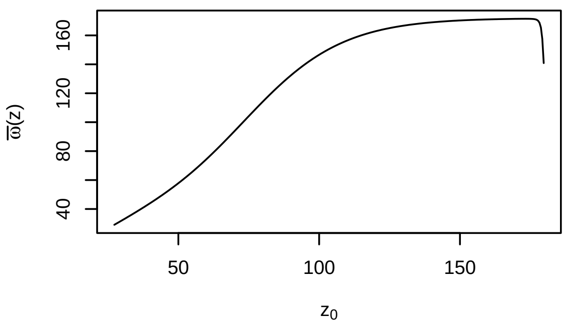

This is an interesting variety of relationships! Sometimes, it increases linearly, whereas other times we get a parabolic relationships. This is a good first pass on an analysis, though subsequent digging would likely reveal quirks in some of these models that we'd need to address more thoroughly. After these exercises, you should have the tools to do just that!

#### Recap

We first showed how to subset PADRINO using the Metadata table, as well as a few others. Next, we demonstrated how to rebuild kernels from the database, as well as basic analyses such as deterministic population growth rates and perturbations. Next we moved into some examples of life cycle properties, such as recruit production and size at death. Along the way, we encountered some issues with data that required us to manipulate the underlying `proto_ipms`. This is far from an exhaustive display of potential analyses. However, this case study should serve as a guide for posing questions and solving issues with PADRINO.
