## Supplementary material for "Rpadrino: an R package to access and use PADRINO, an open access database of Integral Projection Models": Case Study 2

### Combined analyses with Rpadrino, ipmr, and other databases

**Recap** **16**

**Citations** **16**

#### Overview

In this case study, we demonstrate how PADRINO can be used in conjunction with (1) your own unpublished data, and (2) external data repositories.

In many cases, we may wish to combine data that we've collected and not yet published with data from PADRINO. For example, we may wish to compare the vital rates or sensitivities of our (as-yet-unpublished) study system with those from published IPMs on e.g. similar species. We describe how to do this in Section 1 of this case study.

Using PADRINO in conjunction with external data sources presents unique opportunities to address synthetic questions in ecology, evolutionary biology, and conservation. External data sources could (though are not limited to) include climate data, species range distributions, phylogenies, or life tables. We describe how to use PADRINO in conjunction with BIEN in Section 2 of this case study.

#### Combining your own data with PADRINO

In this section, we demonstrate how to combine data from PADRINO with a users own unpublished data. The first part of the code generates two IPMs - these are our "unpublished IPMs". The next section shows how to combine our unpublished IPMs with those stored in PADRINO, and how to perform simple analyses (i.e. calculate population growth rate) across the combined set of IPMs.

#### Creating our own IPMs

The goal of this case study is to show how to combine data, and not necessarily how to use `ipmr`. `ipmr` is extensively documented on the project's website and in the publication describing the package. Therefore, the next few chunks of code assume you have already consulted these resources and will have a reasonable understanding of what's going on. If you have not already consulted these, please do so now.

Our first "homemade" IPM will be a general IPM. For now, we are only going to construct `proto_ipm` objects for each one of these homemade IPMs. Once we have our PADRINO IPMs selected, we will splice everything together and generate actual IPM objects.

```
# Loading Rpadrino automatically loads ipmr, so we do not need to load both.
library(Rpadrino)

# Set up the initial population conditions and parameters.
# These are hypothetical values and do not correspond to any particular
# species.

data_list <- list(
  g_int     = 5.781,
  g_slope   = 0.988,
  g_sd      = 20.55699,
  s_int     = -0.352,
  s_slope   = 0.122,
  s_slope_2 = -0.000213,
  r_r_int   = -11.46,
  r_r_slope = 0.0835,
  r_s_int   = 2.6204,
  r_s_slope = 0.01256,
  r_d_mu    = 5.6655,
  r_d_sd    = 2.0734,
  e_p       = 0.15,
  g_i       = 0.5067,
  sb_surv   = 0.2
)

# Lower bound, upper bound, and number of meshpoints.
L <- 1.02
U <- 624
n <- 500

# Initialize a population vector. The continuous state will have 500 meshpoints,
# and we will pretend there's a seedbank.

init_pop_vec   <- runif(500)
init_seed_bank <- 20

my_general_ipm <- init_ipm(sim_gen = "general", di_dd = "di", det_stoch = "det") %>%
  define_kernel(
    name          = "P",
    formula       = s * g * d_ht,
    family        = "CC",
    g             = dnorm(ht_2, g_mu, g_sd),
    g_mu          = g_int + g_slope * ht_1,
    s             = plogis(s_int + s_slope * ht_1 + s_slope_2 * ht_1^2),
```

```
  data_list     = data_list,
  states        = list(c('ht')),
  uses_par_sets = FALSE,
  evict_cor     = TRUE,
  evict_fun     = truncated_distributions('norm',
                                          'g')
) %>%
define_kernel(
  name          = "go_discrete",
  formula       = r_r * r_s * d_ht,
  family        = 'CD',
  r_r           = plogis(r_r_int + r_r_slope * ht_1),
  r_s           = exp(r_s_int + r_s_slope * ht_1),
  data_list     = data_list,
  states        = list(c('ht', "b")),
  uses_par_sets = FALSE
) %>%
define_kernel(
  name          = "stay_discrete",
  family        = "DD",
  formula       = sb_surv * (1 - g_i),
  data_list     = data_list,
  states        = list(c("b")),
  uses_par_sets = FALSE
) %>%
define_kernel(
  name          = 'leave_discrete',
  formula       = e_p * g_i * r_d * d_ht,
  r_d           = dnorm(ht_2, r_d_mu, r_d_sd),
  family        = 'DC',
  data_list     = data_list,
  states        = list(c('ht', "b")),
  uses_par_sets = FALSE,
  evict_cor     = TRUE,
  evict_fun     = truncated_distributions('norm',
                                          'r_d')
) %>%
define_impl(
  list(
    P              = list(int_rule    = "midpoint",
                          state_start = "ht",
                          state_end   = "ht"),
    go_discrete    = list(int_rule    = "midpoint",
                          state_start = "ht",
                          state_end   = "b"),
    leave_discrete = list(int_rule    = "midpoint",
                          state_start = "b",
                          state_end   = "ht"),
    stay_discrete  = list(int_rule    = "midpoint",
                          state_start = "b",
                          state_end   = "b")
  )
) %>%
```

```
  define_domains(
    ht = c(L, U, n)
  ) %>%
  define_pop_state(
    pop_vectors = list(
      n_ht = init_pop_vec,
      n_b  = init_seed_bank
    )
  )
```

Our next IPM will be a simple one:

```
# Another hypothetical model. These parameters also do not correspond to any
# species.

my_data_list = list(s_int     = -2.2,
                    s_slope   = 0.25,
                    g_int     = 0.2,
                    g_slope   = 0.99,
                    sd_g      = 0.7,
                    r_r_int   = 0.003,
                    r_r_slope = 0.015,
                    r_s_int   = 0.45,
                    r_s_slope = 0.075,
                    mu_fd     = 2,
                    sd_fd     = 0.3)

my_simple_ipm <- init_ipm(sim_gen = "simple",
                          di_dd   = "di",
                          det_stoch = "det") %>%
  define_kernel(
    name      = "P_simple",
    family    = "CC",
    formula   = s * G,
    s         = plogis(s_int + s_slope * dbh_1),
    G         = dnorm(dbh_2, mu_g, sd_g),
    mu_g      = g_int + g_slope * dbh_1,
    data_list = my_data_list,
    states    = list(c('dbh')),
    evict_cor = TRUE,
    evict_fun = truncated_distributions(fun    = 'norm',
                                        target = 'G')
  ) %>%
  define_kernel(
    name      = 'F_simple',
    formula   = r_r * r_s * r_d,
    family    = 'CC',
    r_r       = plogis(r_r_int + r_r_slope * dbh_1),
    r_s       = exp(r_s_int + r_s_slope * dbh_1),
    r_d       = dnorm(dbh_2, mu_fd, sd_fd),
    data_list = my_data_list,
    states    = list(c('dbh')),
    evict_cor = TRUE,
    evict_fun = truncated_distributions(fun    = 'norm',
```

```
                                        target = 'r_d')
  ) %>%
  define_impl(
    make_impl_args_list(
      kernel_names = c("P_simple", "F_simple"),
      int_rule     = rep("midpoint", 2),
      state_start  = rep("dbh", 2),
      state_end    = rep("dbh", 2)
    )
  ) %>%
  define_domains(
    dbh = c(0,
            50,
            100
    )
  ) %>%
  define_pop_state(
    n_dbh = runif(100)
  )

my_ipm_list = list(ipm_1 = my_general_ipm, ipm_2 = my_simple_ipm)
```

#### Combining user-defined and PADRINO-defined IPMs

Next, we will create a list of `proto_ipm` objects from PADRINO, and then put everything together. For simplicity, we will select a small number of plant species. The `pdb` object is contained within the `Rpadrino` package. It is not a complete version of PADRINO. We will use the complete data set in the next section, accessed with `pdb_download()`.

```
data(pdb)

id_index <- c(
  paste0( "aaaa", c(34, 55)),
  paste0("aaa", c(310, 312, 339, 341, 353, 388))
)

small_db <- pdb_subset(pdb, id_index)
```

Next, we need to create a list that holds both the PADRINO IPMs and the ones we created above. After that, we can call `pdb_make_ipm()` on the combined data set, and voila! We have our database IPMs and our own homemade ones.

```
proto_list <- c(
  pdb_make_proto_ipm(small_db),
  my_ipm_list
)
## 'ipm_id' aaa310 has the following notes that require your attention:
## aaa310: 'Geo and time info retrieved from COMPADRE (v.X.X.X.4)'
## 'ipm_id' aaa388 has the following notes that require your attention:
## aaa388: 'Same data as AAA388. State variable Height (Cm)'
```

Great! In that single step, we combined PADRINO IPMs with our own IPMs. Because these are all in the `proto_ipm` format, we do not need to think about technical differences between each type - we can use the exact same toolbox for analyzing both! Let's build the IPM objects and calculate deterministic per-capita growth rates!

```
ipm_list   <- pdb_make_ipm(proto_list)
lambdas    <- lambda(ipm_list)
```

We could now proceed with any further analyses just as we did in the case study 1. Since those types of analyses are already covered by the previous case study, we will move on to combining PADRINO data with information from other databases.

### Extending analyses with other databases

Here, we show how to combine data from PADRINO with data from other external sources. Specifically, we show how to combine with data from COMPADRE MPM database and BIEN , a database containing the spatial distribution and phylogenies of many plant species. We then demonstrate how we can use these combined datasets to address the question: "How do population growth rates vary by the distance of the studied population from the known range edge of that species?"

Given the question we posed above, we need to get range maps for each species and the per-capita growth rate for some populations. For the former, we will use range maps from BIEN.For the latter, we will augment PADRINO with data from COMPADRE. This analysis will not be the most complete - it is intended to demonstrate the steps for combining data, not to make a scientific point. With that in mind, let's dive in!

#### Required packages

BIEN allows users to download range maps programmatically from their database using the BIEN R package. You can install that from CRAN using the chunk below. we will also use `Rcompadre`, `mgcv`, `ggplot2`, `sf`, and `dplyr` to work with the data, so you'll need to install those as well.

```
install.packages(c("BIEN", "ggplot2", "sf", "dplyr", "Rcompadre", "mgcv"))
```

After that, we have to load them:

```
library(BIEN)
library(ggplot2)
library(sf)
library(dplyr)
library(Rcompadre)
library(Rpadrino)
library(mgcv)
```

#### Data identification

BIEN allows us to programmatically query the database and retrieve all species names for which there is a range map. we will load that, then load COMPADRE and PADRINO, and see how much overlap there is.

```
bien_rng_spps <- BIEN_ranges_list()
pdb           <- pdb_download(save = FALSE)
cdb           <- cdb_fetch("compadre")
```

```
## This is COMPADRE version 6.21.8.0 (release date Aug_20_2021)
## See user agreement at https://compadre-db.org/Help/UserAgreement
## See how to cite at https://compadre-db.org/Help/HowToCite
```

```r
# Insert an underscore to make sure name format matches between COMPADRE,
# PADRINO, and BIEN

cdb_spp <- gsub(" ", "_", cdb$SpeciesAccepted)

pdb_spp <- pdb$Metadata$species_accepted
```

Nice! We have 480 overlapping species between COMPADRE/PADRINO and BIEN's range maps. This next chunk determines which species from PADRINO and COMPADRE have range maps available in BIEN:

```r
all_spp <- unique(c(cdb_spp, pdb_spp))
pos_spp <- all_spp[all_spp %in% bien_rng_spps$species]

pdb_rng_spp <- unique(pdb_spp[pdb_spp %in% pos_spp])
cdb_rng_spp <- unique(cdb_spp[cdb_spp %in% pos_spp])
```

#### Subsetting

We probably should not use all of these, as those calculations would take quite some time for a tutorial, so we will select a subset. we will take the species for which the demographic data are from North America. For PADRINO, we need to find their `ipm_ids`, and then pass those into `pdb_subset()`. For COMPADRE, we can just use `dplyr` verbs as if we were working with a `data.frame`.

```r
# First, we will create a vector of ipm_id's which meet the following requirements:
# 1. They have range maps in BIEN (species_accepted %in% pdb_rng_spp)
# 2. The model is from data collected in North America (continent == "n_america")
# 3. The data are from unmanipulated populations (treatment == "Unmanipulated")

pdb_ids     <- pdb$Metadata$ipm_id[pdb$Metadata$species_accepted %in% pdb_rng_spp &
                                   pdb$Metadata$continent == "n_america" &
                                   pdb$Metadata$treatment == "Unmanipulated"]

use_pdb     <- pdb_subset(pdb, pdb_ids)

# For COMPADRE, we have to first replace "_" in the species names with a space.
# Then we can use filter() syntax to subset COMPADRE to the species we want.

cdb_rng_spp_f <- gsub("_", " ", cdb_rng_spp)

use_cdb <- filter(cdb,
                  SpeciesAccepted %in% cdb_rng_spp_f &
                  Continent == "N America" &
                  MatrixTreatment == "Unmanipulated")
```

#### Check data quality

PADRINO data is validated before it is uploaded to ensure the IPM behaves as the published version behaves. There are additional checks you might want to perform on your own, and those depend on the subsequent anaylsis. Case study 1 shows an example of a singular kernel creating some biologically impossible results. However, there are not built-in functions in `Rpadrino` yet to assist with this. Therefore, it is usually a good idea to check the original publications just to be sure there are not caveats to the model that the authors have raised. We can find the citations using `pdb_citation()` and `pdb_report()`. `pdb_citation()` returns a character vector of citations in APA style, whereas `pdb_report()` generates an RMarkdown report based on the information in the database.

```
cites <- pdb_citations(use_pdb)

pdb_report(use_pdb)
```

we will also want to check COMPADRE for some common data issues using the `cdb_flag()` function. This is documented much more thoroughly in the `Rcompadre` package website. For simplicity, we will just use ones which do not raise any flags, as fixing issues with COMPADRE data is beyond the scope of this case study. Furthermore, we will subset out the mean matrices, as we want to work with individual transitions.

```
cdb_f <- cdb_flag(use_cdb)

use_cdb <- filter(cdb_f, !check_NA_A & !check_NA_U & !check_NA_F & !check_NA_C&
                         !check_zero_U & !check_singular_U & check_component_sum &
                         check_ergodic & check_irreducible & check_primitive &
                         check_surv_gte_1 & MatrixComposite == "Individual")
```

#### Data transformation

Next, we need to do a bit of data wrangling. From PADRINO, we only need the `ipm_id` and species names for plotting and analyzing, so we will just grab those from the metadata table. We're going to create an `sf` object for this data using the coordinates stored in the `"lat"` and `"lon"` columns of the metadata. `sf` provides a standardized interface for dealing with multiple types of spatial data, and also plays nicely with `dplyr`, which makes managing data much easier. The `st_as_sf()` function handles the conversion for us.

```
# Create a standard data.frame with ipm_id, species, lat+lon data from PADRINO

temp_coords <- use_pdb$Metadata %>%
  select(ipm_id, species_accepted, lat, lon)

# This next bit does the following:
# 1. Creates a data.frame from COMPADRE data with the same columns as PADRINO.
# 2. Changes the names so they match the PADRINO version.
# 3. Combines the COMPADRE and PADRINO versions.
# 4. Eliminates studies that do not have complete latitude/longitude information.

temp_db <- use_cdb@data %>%
  select(MatrixID, SpeciesAccepted, Lat, Lon) %>%
  setNames(names(temp_coords)) %>%
  rbind(temp_coords) %>%
  .[complete.cases(.), ]

# Finally, create an 'sf' object with the combined coordinates from COMPADRE and
# PADRINO

study_coords <- st_as_sf(temp_db,
                         coords = c("lon", "lat"),
                         crs = "WGS84")
```

#### Querying BIEN

Now that we have our final species list, we're going to download the range maps for each species using the `BIEN_ranges_load_species()` function, and then convert that into an `sf` object which will make subsequent analysis and plotting easier.

After converting the range maps to an `sf` object, we also need to create a different version of the polygons that are a set of lines representing the edges. This will allow us to quickly calculate the distance between our study points and the edge of the range. `st_cast()` handles this conversion for us.

```r
# First, convert species names back to spaced version without "_"s. while the
# data.frame of species names we downloaded from BIEN before uses an "_" in
# species names, the `BIEN_ranges_load_species()` function expects names without
# underscores!

study_coords$species_accepted <- gsub("_", " ", study_coords$species_accepted)

# The next piece:
# 1. Downloads range maps
# 2. Converts each range map into an 'sf' object
# 3. Resolves issues that arise in the conversion process (e.g. self intersections)
# 4. Sorts the resulting 'sf' object alphabetically on species

rng_maps <- BIEN_ranges_load_species(study_coords$species_accepted) %>%
  st_as_sf() %>%
  st_make_valid() %>%
  arrange(species)

# We need to create a copy of each range map that, rather than a polygon format,
# is a line format. This enables us to use functions to compute distance to edge
# much more easily.

line_maps <- st_cast(rng_maps, "MULTILINESTRING")

# Put the "_"'s back into study_coords so that we can match all names later on.

study_coords$species_accepted <- gsub(" ", "_", study_coords$species_accepted)

# Finally, sort study_coords alphabetically as well. Now, the species_accepted
# column should be identical to the species column in rng_maps. This will be
# important in the next step.

study_coords <- arrange(study_coords, species_accepted)  %>%
  .[!duplicated(.$species_accepted), ]
```

#### Compute distance from edges

Ok, we're finally ready to compute the distance from each study site to the range edge. We're going to use the `st_distance()` function for this. This finds the minimum distance between the first and second arguments and computes a matrix for all possible combinations. It will ignore the fact that sometimes the closest edge is an ocean (which our species cannot grow in). However, working out how to improve that calculation is a problem for another day!

We start by extracting a distance matrix and taking the diagonal. The diagonal represents the shortest distance between our species study site and the edge of the polygon of its range map (NB: This only works because we sorted each object alphabetically ahead of time!). Next, we add in the species name information and set the data frame's names to something useful. Finally, we will convert the distances to kilometers.

```
# Quickly check to make sure all of our species line up positionally.
# If not, we'd need to make sure they do, otherwise it will be difficult
# to extract the distances from the distance matrix we are about to compute!

stopifnot(all(line_maps$species == study_coords$species_accepted))

# This next piece does:
# 1. Computes distances between all pairs of points and line objects
# 2. Extracts the diagonal of the distance matrix. This represents the distance
# from a species' study site to the edge of its range (again, only because
# we sorted study_coords and line_maps alphabetically).
# 3. converts this information to a data.frame
# 4. Adds the species names to that data.frame
# 5. Sets column names for that data.frame
# 6. Converts the distance to km

dist_from_edge <- st_distance(study_coords, line_maps) %>%
  diag() %>%
  data.frame() %>%
  cbind(study_coords$species_accepted, .) %>%
  setNames(c("species", "distance_in_meters")) %>%
  mutate(
    distance_in_km = round(as.numeric(distance_in_meters) / 1e3, 2)
  )
```

#### Visualize our dataset

we will plot our range maps with the study sites overlaid on them using `ggplot2`. `ggplot2` has built in `geoms` designed to handle `sf` objects, which will make our lives much easier!

```
world     <- map_data("world")
n_america <- filter(world, region %in% c("USA", "Canada", "Mexico"))

ggplot(rng_maps) +
  geom_sf(aes(fill = species), alpha = 0.4) +
  geom_polygon(data = n_america, aes(x = long, y = lat, group = group),
               inherit.aes = FALSE,
               color = "grey70",
               fill = NA) +
  geom_sf(data  = study_coords) +
  coord_sf(xlim = c(-140, -55),
           ylim = c(15, 50)) +
  theme_bw() +
  theme(
    legend.position = "none"
  )
```

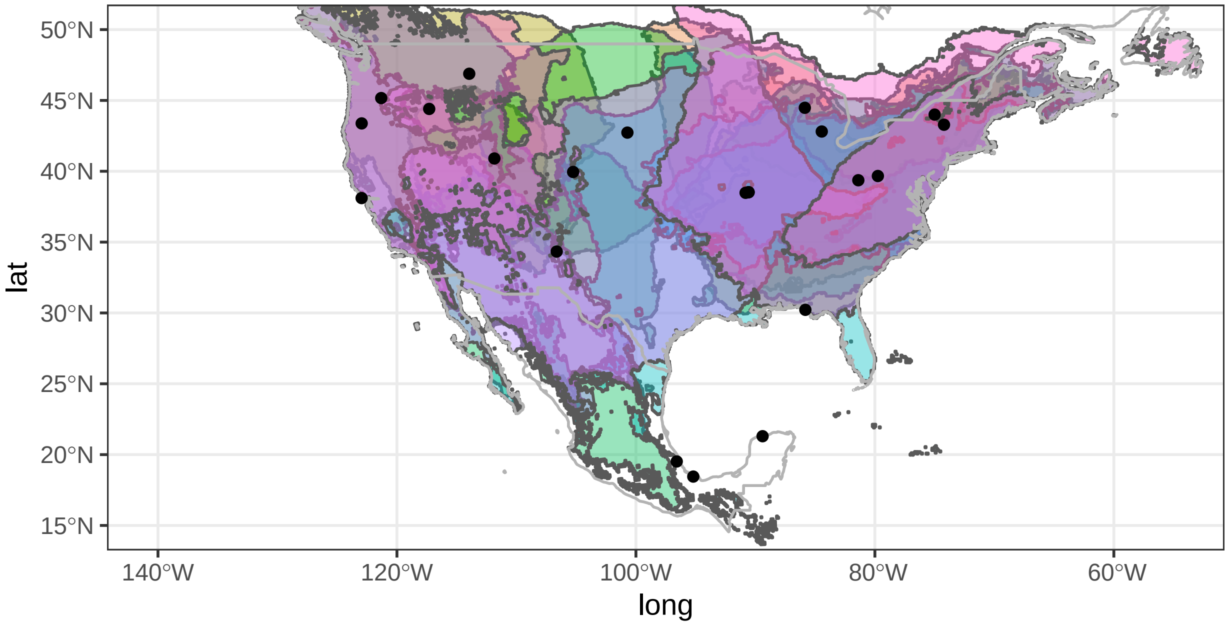

Already, we can see that our range maps do not perfectly align with the COMPADRE and PADRINO population coordinates. We can check and see which study populations are actually contained by their range map like so:

```
# Notice that now we are using the POLYGONS object (rng_maps) as opposed to the
# to the LINESTRING version (line_maps).

covered_ind <- st_covered_by(study_coords,
                             st_make_valid(rng_maps),
                             sparse = FALSE) %>%
  diag()

# Print studies not covered by BIEN range map
study_coords[!covered_ind , ]
```

```
## Simple feature collection with 6 features and 2 fields
## Geometry type: POINT
## Dimension:     XY
## Bounding box:  xmin: -122.9567 ymin: 18.45 xmax: -85.82 ymax: 45.16667
## Geodetic CRS:  WGS 84
## # A tibble: 6 x 3
##   ipm_id species_accepted                    geometry
##   <chr>  <chr>                            <POINT [°]>
## 1 241875 Astragalus_tyghensis   (-121.3167 45.16667)
## 2 242052 Calathea_ovandensis           (-95.2 18.45)
## 3 239708 Euphorbia_telephioides    (-85.82 30.21972)
## 4 244332 Lupinus_tidestromii    (-122.9567 38.10861)
```

```
## 5 247007 Mammillaria_gaumeri            (-89.4 21.3)
## 6 241182 Tillandsia_deppeana    (-96.58333 19.51667)
```

These are COMPADRE matrices. Rather than try to figure out what's going on, we will just drop those out of our analysis.

```
study_coords <- study_coords[covered_ind, ]
```

#### Compute lambdas for each type of model

Great! The next step is to generate and then join our lambda values with the distance information. This is a two-step process. First, we will build our PADRINO IPMs:

```
# Extract PADRINO IDs - we do not want to give COMPADRE ones to PADRINO machinery!

pdb_ids <- study_coords$ipm_id[study_coords$ipm_id %in% pdb$Metadata$ipm_id]

# Construct the proto_ipm list
proto_list <- pdb_make_proto_ipm(
  use_pdb,
  pdb_ids
)
```

```
## 'ipm_id' aaa310 has the following notes that require your attention:
## aaa310: 'Geo and time info retrieved from COMPADRE (v.X.X.X.4)'

## 'ipm_id' aaa329 has the following notes that require your attention:
## aaa329: 'Based on IPM from Rose Ecology 2005; The GPS coordinates were approximated
## to the closest geographic location described in the reference'

## 'ipm_id' aaa385 has the following notes that require your attention:
## aaa385: 'Same data as AAA385. State variable Height (Cm)'
```

```
# Construct the IPMs

ipm_list <- pdb_make_ipm(proto_list)

# Some IPMs may have many values for lambda, because they were constructed from
# vital rate models that have time varying parameters (e.g. random effects for
# year). we will need to account for this. We need to convert those from a list to
# a data.frame for modeling, and need to keep track of which lambda belongs to
# which ID. The loop below will correctly format this.

lambdas <- lambda(ipm_list, type_lambda = "last")

temp <- data.frame(ipm_id = NA,
                   lambda = NA)

for(i in seq_along(lambdas)) {

  # Create a temporary object to store lambda values and ipm_id's. Each lambda
  # value will have its own row, with the corresponding ipm_id next to it. This
  # will help us track which value belongs to which model.

  temp_2 <- data.frame(ipm_id = names(lambdas)[i],
                       lambda = lambdas[[i]])
```

```r
  # I don't normally recommend using rbind in a 'for' loop, but there aren't many
  # iterations here, so we will not worry about the memory footprint
  temp <- rbind(temp, temp_2)

}

# Remove the dummy row of NAs

temp <- temp[-1, ]
```

Next, we will get our COMPADRE lambdas, and stick them back in with the PADRINO lambdas.

```r
use_cdb <- filter(use_cdb, MatrixID %in% study_coords$ipm_id)

matAs   <- matA(use_cdb)

use_cdb@data$lambda <- vapply(matAs,
                              function(x) Re(eigen(x)$values[1]),
                              numeric(1L))

cdb_lambda <- use_cdb@data %>%
  select(MatrixID, lambda) %>%
  setNames(c("ipm_id", "lambda"))

all_lambdas <- rbind(temp, cdb_lambda)
```

Finally, we need to join lambda values with coordinate data set to recover the species names, and then use those to join with the distance from edge object. Once that's done, we can plot everything!

```r
all_lambdas <- left_join(all_lambdas, study_coords, by = "ipm_id") %>%
  select(-geometry)

all_data <- left_join(all_lambdas, dist_from_edge,
                      by = c("species_accepted" = "species"))
```

#### Regression modelling

We're ready to plot and analyze the data. GAMs (Wood 2011) are a great way to spot general trends in data, so we will use those.

```r
lambda_by_dist <- gam(lambda ~ s(distance_in_km, bs = "cs"),
                      data = all_data,
                      family = Gamma(link = "identity"))

summary(lambda_by_dist)
```

```
##
## Family: Gamma
## Link function: identity
##
## Formula:
## lambda ~ s(distance_in_km, bs = "cs")
##
```

```
## Parametric coefficients:
##             Estimate Std. Error t value Pr(>|t|)
## (Intercept)  1.13381    0.04843   23.41 9.22e-16 ***
## ---
## Signif. codes:  0 '***' 0.001 '**' 0.01 '*' 0.05 '.' 0.1 ' ' 1
##
## Approximate significance of smooth terms:
##                     edf Ref.df     F p-value
## s(distance_in_km) 6.463      9 1.305   0.135
##
## R-sq.(adj) =  0.197   Deviance explained = 47.5%
## GCV = 0.059745  Scale est. = 0.048724  n = 27
```

```r
plot(lambda_by_dist)
```

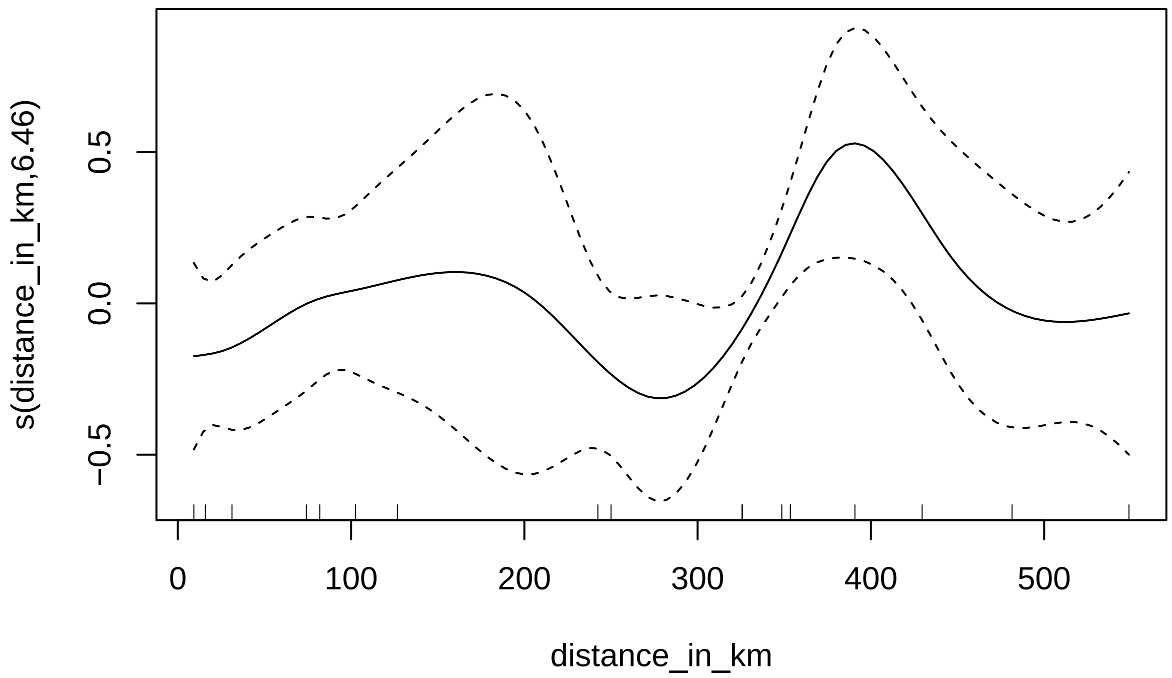

```r
preds <- cbind(data.frame(predict(lambda_by_dist,
                                  data.frame(distance_in_km = seq(0, 550, 1)),
                                  type = "response",
                                  se.fit = TRUE)),
               x = seq(0, 550, 1)) %>%
  mutate(upper = fit + se.fit * 1.96,
         lower = fit - se.fit * 1.96)

ggplot(all_data, aes(x = distance_in_km, y = lambda)) +
  geom_point() +
```

```
geom_smooth(method = "glm",
            formula = y ~ x,
            method.args = list(family = Gamma("identity")),
            color = "red",
            linetype = "dotted") +
geom_line(data = preds,
          aes(x = x, y = fit),
          inherit.aes = FALSE,
          color = "blue",
          size = 1.2) +
geom_ribbon(data = preds,
            aes(x = x, ymin = lower, ymax = upper),
            color = "grey",
            alpha = 0.4,
            inherit.aes = FALSE) +
theme_bw() +
xlab("Distance from range edge (km)") +
ylab(expression(lambda))
```

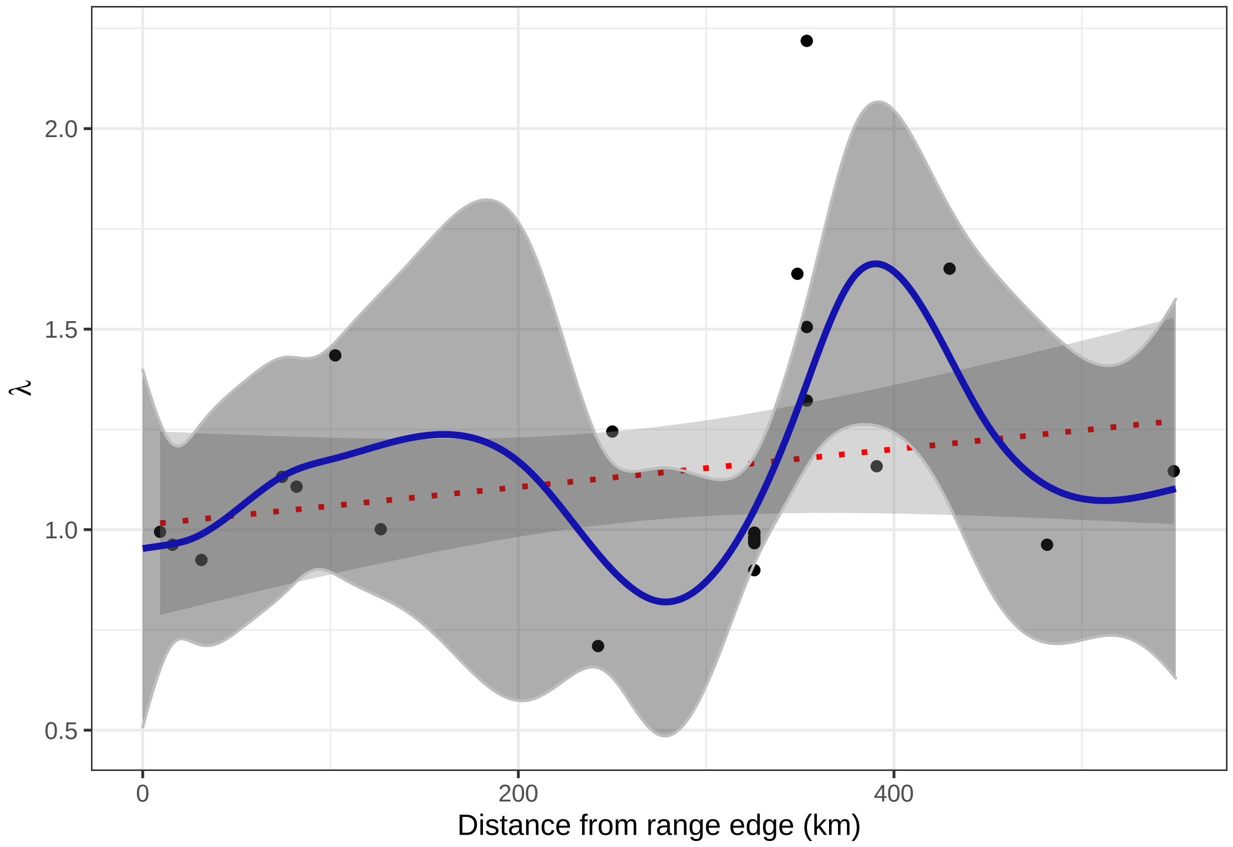

There is a positive trend in range centrality and species performance (red line), and the GAM is likely overfit (blue line). There is a lot of residual variance, and we can certainly find better ways to model this phenomenon, but this is a good start for an exploratory analysis. We will leave the further analyses as an exercise to you!

### Recap

We have shown how to combine PADRINO data with user-defined IPMs as well as join it with information from other databases. This is hardly a comprehensive overview of PADRINO's applications - there are many other uses and databases one could combine PADRINO with. It is our hope that this and the previous case study provide a general guide to the considerations and steps one needs to take when using this data!
