## Supplementary material for "Rpadrino: an R package to access and use PADRINO, an open access database of Integral Projection Models": Electronic Supplementary Info

**Technical overview of PADRINO** **2**

### Introduction to PADRINO

PADRINO v0.0.1 consists of 10 tables (Table 1, Figure S1). In this first version, PADRINO currently contains 280 IPMs from 40 peer-reviewed publications that consider 14 animal and 26 species (Table 1 main text). However, we highlight that PADRINO is under active development, and we continue to digitize studies for release in future versions. These tables form a database, with tables linked using a common column across all tables: `ipm_id`. The scope of each `ipm_id` is determined by the way that an IPM is parameterized. IPMs that characterize the same species across, for example, many years or sites, with the same functional form, are included under a single `ipm_id`. For instance, a growth model that includes a random intercept for different years could be used to generate many unique projection kernels. These are stored under a single `ipm_id` because the functional form of the IPM is identical for each year, and only the parameter values change. One exception to this grouping rule is when the sites (*i.e.* where the raw data are reported to have come from) are far enough apart that separate sets of GPS coordinates are used to describe them. These IPMs are split into separate `ipm_id`s so that the spatial distinctions are preserved, which facilitates matching PADRINO data with, for example, gridded environmental data (*e.g.* Compagnoni et al. 2021b, Case Study 2).

Finally, there are two important details potential users should be aware of. The first detail is that PADRINO provides IPMs *as they are published following peer review.* We do not alter these IPMs when digitizing them, except to correct typographical errors that may have found their way into the peer-reviewed publication. The second detail is that PADRINO does not store any raw data used to create the IPMs. Users should be aware of these, and we encourage all users to consult and cite the original publications of each IPM before including it in an analysis.

#### The Digitization Process

The IPM digitization process begins when a peer-reviewed paper containing an IPM is published. We have set alerts for the following keyword searches: "Integral Projection Model OR IPM OR sensitivit* OR elasticit* OR Vital rate OR LTRE". This automatic weekly search is run on Google Scholar and Scopus, and resulting hits are examined manually to find publications that contain an IPM. Once a paper containing an IPM is identified, we extract five types of metadata: taxonomic information (*e.g.* species names, functional groups), publication information (*e.g.* authors, complete citation, year of publication), temporal metadata (*e.g.* study duration, data collection beginning and ending months and years), spatial metadata (*e.g.* latitude/longitude, ecoregion), and model specific metadata (*e.g.* experimental treatments applied, density-(in)dependent). Table S1 contains a complete description of the metadata table in PADRINO.

Following the metadata digitization, we extract functional forms of each sub-kernel, vital rate function, and how the environment varies (if applicable). The functional forms of each component of the model are expressed in the syntax introduced in the main text. Finally, we extract all of the parameter values, as well as information on the range of values each trait can take on and how they are numerically approximated (*i.e.* integration rules). The parameter values and integration information are then substituted for symbol names when the user requests a built model. For example, in *Rpadrino*, the `Norm(mu_g, sd_g)` from the main text would be translated to `dnorm(z_2, mu_g, sd_g)`.

Often times, not all of the required information is present in the publication or its supplementary materials. Therefore, we often contact authors to request the required information and/or ask for clarification. We also extract a target value for the data validation step (see next section), so that we can ensure that released data really does replicate the published IPM. A complete guide to our digitization process and documentation of the database syntax is publicly available on PADRINO's webpage (https://padrinoDB.github.io/Padrino/).

##### Data Validation and Reproducibility

The PADRINO IPM Database has automated testing built into the data release process. All IPMs are checked to ensure they recover the behavior of the published version prior to release. In most cases, validation consists of reproducing the kernel-specific asymptotic population growth rate ($\lambda$) to within $\pm$ 0.03 of the published $\lambda$ value in the source publication. It is worth noting that this margin of error is considerably lower than the uncertainty that arises from fitting statistical models to the raw data used in the IPM (*e.g.* Clark 2003), and so it should be acceptable for almost any application. For stochastic models with continuously varying environments, it is often not computationally feasible to re-run the IPM for 10-50,000 iterations since they are time consuming to run and there are many in PADRINO. Thus, we manually check for shorter term behavior that is similar to published dynamics (*e.g.* stochastic population growth rate ($\lambda_s$) after 1000 iterations). For publications where population growth rates are not available, we manually examine the publication and check the model digitized in PADRINO against some reported behavior (*e.g.* generation time). A given IPM can only enter a scheduled database release if it is explicitly flagged by a digitizer as validated, or if it passes its automated test. The manual testing functionality is contained in the open source *R* package *pdbDigitUtils* (available on GitHub (https://github.com/padrinoDB/pdbDigitUtils)), and PADRINO's build scripts are in the project's GitHub repository (https://github.com/padrinoDB/Padrino/tree/main/R).

##### Challenges

Digitizing IPMs into the PADRINO IPM Database is not without issues. First, it is often the case that the complete form of the IPM is not reported: approximately 80% of papers we have examined thus far fall into this category. Many studies may report the general form of the model (*e.g.* $n(z', t+1) = \int_L^U K(z', z)n(z, t)dz$), but do not then report the functional forms of the sub-kernels or vital rates. Without the functional form of all vital rates and sub-kernels, it is impossible to reproduce the IPM. Second, some parameter values may be missing from the main text or supplementary materials - common culprits are terms for the variation of the growth/fecundity kernels, number of meshpoints, and integration bounds (i.e. $L, U$ in Eq 1). The authors of this paper have been guilty of this, as well as other sins of omission, in their own IPM publications. The intent here is not to alienate other authors, but offer a gentle reminder that reporting all parameter values and functional forms can go a long way towards making their science reusable and extensible. Reproducible science can often bring great benefit to the original authors as well as the broader community (Kousta et al. 2019).

#### Technical overview of PADRINO

PADRINO is structured such that each model gets one row for the Metadata table, and an arbitrary number of rows for every table after that. Some models may have 0, 1, or many rows for some of these tables. Information for each model is linked across tables by the `ipm_id` column. Complete descriptions of each column are provided here.

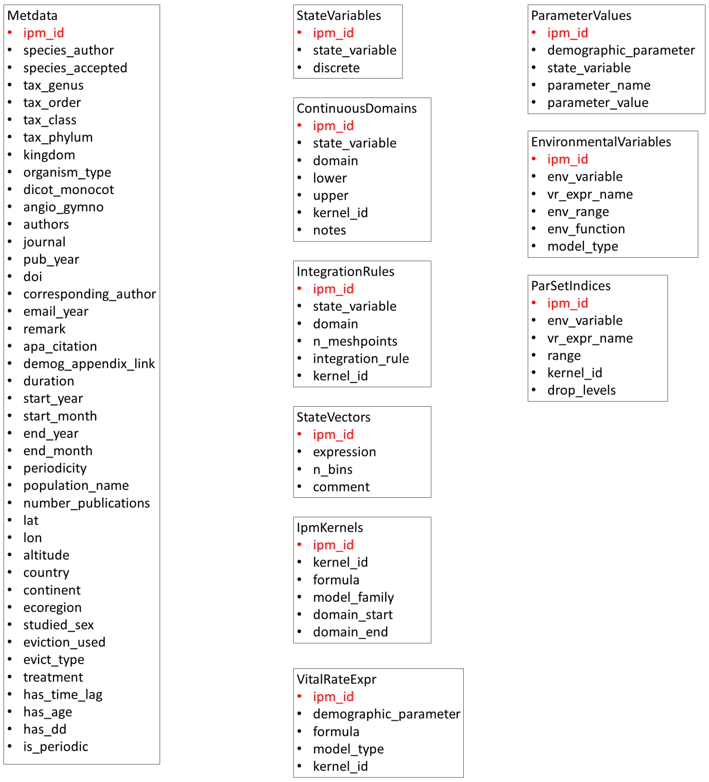

When a user calls `pdb_make_proto_ipm()` and specifies `ipm_ids`, the function loops over the specified IDs subsetting the database to each single one. It then calls `.make_proto()`, which first translates each IPM component from PADRINO syntax into `ipmr` syntax, then calls `define_*` functions from `ipmr` to generate a `proto_ipm`. If there is more than one ID requested, then `pdb_make_proto_ipm()` repeats the process as many times as requested to generate a list of `proto_ipm`s. This list can be passed to `pdb_make_ipm()`, `pdb_make_ipm()` is a thin wrapper around `ipmr`'s `make_ipm()`, and allows for different sets of additional arguments to be passed to each individual IPM build process.

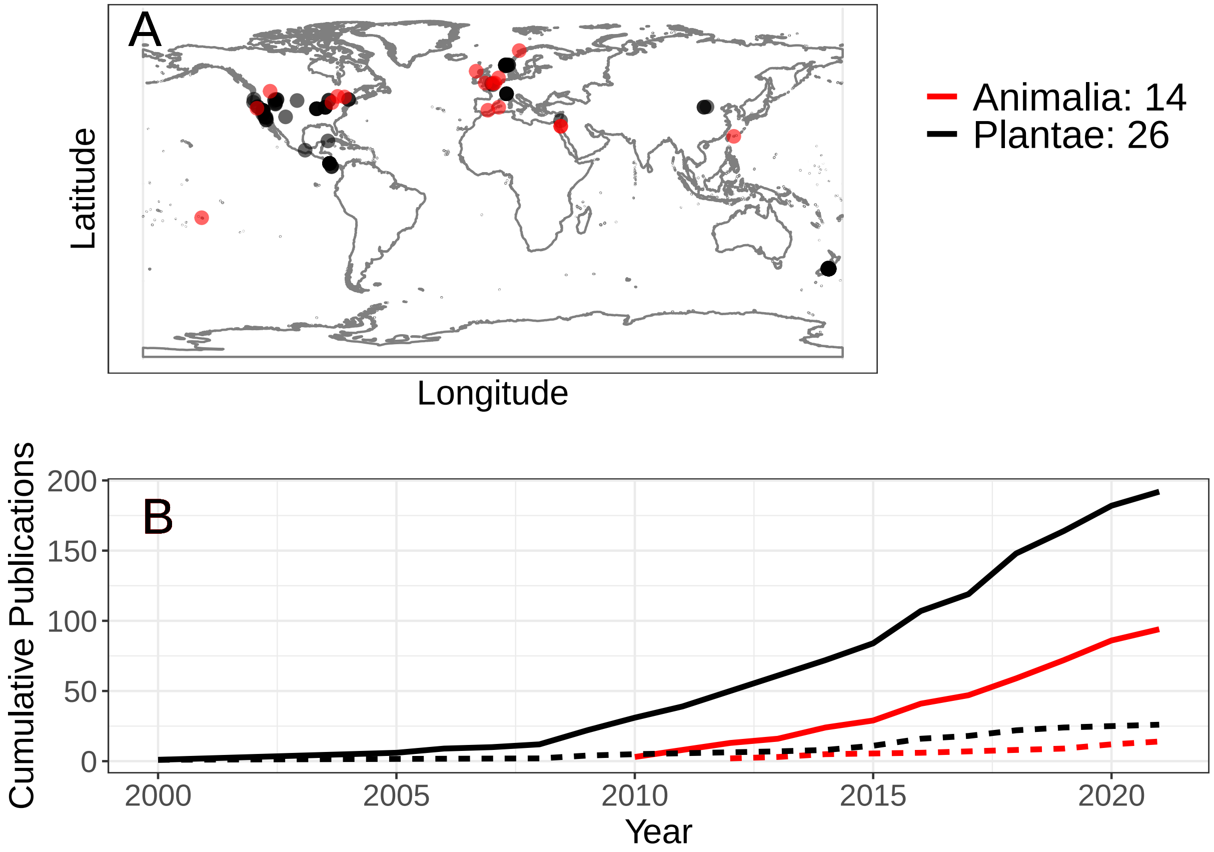

**Figure S1**: The geographic and temporal coverage of studies in the PADRINO IPM Database. (A) Geographic distribution of publications currently contained in PADRINO (i.e. studies from Table 1). (B) Cumulative number of publications found by our search criteria by year (solid lines), and the number that are in the released version of PADRINO (dashed lines). Future releases will include those that we have found, but are not yet completely digitized (i.e. those represented by solid lines, but not yet included in the dashed lines). See the Supplementary Data for a complete list of IPM publications.

Table S1: Summaries of the information contained in each table of the PADRINO database. A complete guide to each column in each table is available on the project's webpage in the form of the guide provided to digitizers (there are too many columns to provide the information here).

| Table | Description |
|---|---|
| Metadata | This table contains metadata for each IPM. This is organized into taxonomic information (full taxonomy plus functional group information), publication information (citation, authorship, source), data collection information (study period/duration, GPS coordinates, ecoregion), and model specific information (studied sexes, eviction corrections, treatments applied, and model implementation details). See Table S2 for more information on these columns. |
| State Variables | This table contains the names of the state variables used in the model and whether or not they are discrete or continuously distributed. |
| Continuous States | This table contains names and ranges for each continuously distributed state variable in the model, as well as which kernels they apply to (kernels are the P(z',z), F(z',z), and C(z',z) in Main Text's Eq 1). |
| Integration Rules | This table contains information on how each continuous state variable is numerically approximated in the model (i.e. number of meshpoints, which integration rule was used). |
| Population Trait Distributions | This table contains the names of the population trait distributions used in the model (n(z,t) and n(z',t+1) in Main Text's Eq. 1). |
| IPM Sub-kernels | This table contains the functional forms of each sub-kernel in the IPM (_e.g._ P(z',z) in Main Text's Eq 1 becomes 'P = s * G'), and information on which traits it acts on and creates. This table makes use of ipmr's [parameter set index notation](https://levisc8.github.io/ipmr/articles/index-notation.html) to concisely represent models which may produce many kernels. |
| Vital Rate Functions | This table contains the functional forms of each vital rate in the IPM (e.g. 'mu__g = int__g + slope__g * z__1'). This table makes use of ipmr's [parameter set index notation](https://levisc8.github.io/ipmr/articles/index-notation.html) to concisely represent models which may produce many kernels. |
| Parameter Values | This table contains the names and values of each parameter in the model, with the exception of parameters that are associated with continuous environmental variation. |
| Continuous Environmental Variation | This table contains parameter values and functional forms of any continuously varying environmental conditions (e.g. yearly variation in precipitation and/or temperature). Any model that contains information in this table is considered stochastic by default, as these variables must be sampled at least once to construct a model with Rpadrino. |
| Parameter Set Indices | This table contains the parameter set indices. These are substituted into the IPM kernels and vital rate expressions when a model is built, so that a single symbolic expression can represent an arbitrary number of realized expression. For example, the vital rate expression 'mu__g__yr = g__int__yr + g__slope * z__1' can be used to represent a range of years for a model with year-specific intercepts. This table contains values substituted in for '__yr' across the model. See the [ipmr vignette on Index Notation](https://levisc8.github.io/ipmr/articles/index-notation.html) for more details. |

Table S2: All columns contained in the Metadata table.

| Concept | Column Name | Description |
|---|---|---|
| | ipm_id | Unique ID for each model. |
| Taxonomy | species_author | The Latin species name used by the authors of the paper. |
| | species_accepted | The Latin species name accepted by Catalogue of Life. |
| | tax_genus | The genus name accepted by Catalogue of Life. |
| | tax_family | The family name accepted by Catalogue of Life. |
| | tax_order | The order name accepted by Catalogue of Life. |
| | tax_class | The class name accepted by Catalogue of Life. |
| | tax_phylum | The phylum name accepted by Catalogue of Life. |
| | kingdom | The kingdom name accepted by Catalogue of Life. |
| | organism_type | General functional type of the species (_e.g._ annual, fern, mammal, reptile). |
| | dicot_monocot | If a plant species, whether the species is a dicot or a monocot. |
| | angio_gymno | If a plant species, whether the species is an angiosperm, gymnosperm, or neither. |
| Source | authors | All of a study authors' last names, separated by ';'. |
| | journal | Abbreviated journal name (www.abbreviations.com/jas.php), or 'PhD', 'MSc' if a thesis. |
| | pub_year | The year of publication. |
| | doi | Digital object identifier and/or ISBN (if available). |
| | corresponding_author | The name of the corresponding author on the paper. |
| | email_year | The email address of the corresponding author and the year it was extracted (some email addresses may be defunct now). |
| | remark | Additional remarks from the digitizer regarding the publication, if any. |
| | apa_citation | The full APA citation for the source. |
| | demog_appendix_link | The URL for the Supplementary information containing additional model details, if available. |
| Temporal Metadata | duration | The duration of the study, defined 'study_end - study_start + 1'. Does not consider skipped years. |
| | start_year | The year demographic data collection began. |
| | start_month | The month demographic data collection began. |
| | end_year | The year demographic data collection ended. |
| | end_month | The month demographic data collection ended. |
| | periodicity | Frequency of the model (1: annual transition, 2: semi-annual transition, 0.2: 5 year transition). |
| Spatial Metadata | population_name | The name of the population given in the data source. |
| | number_populations | The number of populations that a given model describes. |
| | lat | The decimal latitude of the population. |
| | lon | The decimal longitude of the population. |
| | altitude | The altitude of the population above sea level, obtained either from the publication or Google Earth. |
| | country | The ISO3 code for the country or countries in which the data were collected. |
| | continent | The continent or continents on which the data were collected. |

Table S2: All columns contained in the Metadata table. *(continued)*

| Concept | Column Name | Description |
|---|---|---|
| | ecoregion | The terrestrial or aquatic ecoregion corresponding to the [World Wildlife Fund](https://www.worldwildlife.org/biomes) classification. If data are from a controlled setting (greenhouse, lab), denoted with 'LAB'. |
| Model-specific metadata | studied_sex | Sexes used to construct the model. |
| | eviction_used | Whether or not the authors explicitly state that they corrected for eviction (see Williams et al. 2012). |
| | evict_type | If the authors did correct for eviction, then the type of correction that was applied. Current options are 'stretched_domain', 'truncated_distributions', and 'disctrete_extrema'. |
| | treatment | A description of any experimental treatment applied to the population. |
| | has_time_lag | Whether or not the model contains a time lagged vital rate/kernel. |
| | has_age | Whether or not the model has age structure in addition to other continuous state variables. |
| | has_dd | Whether or not the model is density dependent. |
| | is_periodic | Whether or not the model is periodic. |
